# RAD51-mediated R-loop formation acts to repair transcription-associated DNA breaks driving antigenic variation in *Trypanosoma brucei*

**DOI:** 10.1101/2023.05.11.540369

**Authors:** Mark John Girasol, Marija Krasilnikova, Catarina A. Marques, Jeziel D. Damasceno, Craig Lapsley, Leandro Lemgruber, Richard Burchmore, Dario Beraldi, Ross Carruthers, Emma M. Briggs, Richard McCulloch

**Author notes:** Correspondence to and/or.

## Abstract

RNA-DNA hybrids are epigenetic features of all genomes that intersect with many processes, including transcription, telomere homeostasis and centromere function. Increasing evidence suggests RNA-DNA hybrids can provide two conflicting roles in the maintenance and transmission of genomes: they can be the triggers of DNA damage, leading to genome change, or can aid the DNA repair processes needed to respond to DNA lesions. Evasion of host immunity by African trypanosomes, such as *Trypanosoma brucei*, relies on targeted recombination of silent Variant Surface Glycoprotein (*VSG*) genes into a specialised telomeric locus that directs transcription of just one *VSG* from thousands. How such *VSG* recombination is targeted and initiated is unclear. Here, we show that a key enzyme of *T. brucei* homologous recombination, RAD51, interacts with RNA-DNA hybrids. In addition, we show that RNA-DNA hybrids display a genome-wide co-localisation with DNA breaks, and that this relationship is impaired by mutation of RAD51. Finally, we show that RAD51 acts to repair highly abundant, localised DNA breaks at the single transcribed *VSG*, and that mutation of RAD51 alters RNA-DNA hybrid abundance both around the transcribed *VSG* and across the silent *VSG* archive. This work reveals a widespread, generalised role for RNA-DNA hybrids in directing RAD51 activity during recombination and uncovers a specialised application of this interplay during targeted DNA break repair needed for the critical *T. brucei* immune evasion reaction of antigenic variation.

## Introduction

Rapid change in genetic content or organisation, either in localised regions or across the genome, is integral to the life of many organisms. Examples of localised genome variation in eukaryotes include yeast mating type switching^1, 2^ and immunity gene rearrangements to allow the expression of T cell receptors and B cell antibodies^3^. Variation encompassing larger parts of genomes include chromosome fragmentation during development of ciliates^4^and chromosome and gene copy number variation during the life cycle of *Leishmania* ^5, 6^. A myriad of strategies induces such changes, including the generation of DNA breaks by sequence-specific endonucleases, locus-directed replication stalling, and base modification. In most cases, the DNA lesions generated are repaired to effect genetic change, normally by harnessing generalised DNA repair pathways, including homologous recombination (HR), non-homologous end-joining and microhomology-mediated end-joining. Antigenic variation is a widespread immune evasion strategy in which surface antigens are periodically switched to escape elimination of the infecting pathogen by host adaptive immunity^7^. Though some such antigen switches can be catalysed by transcriptional control mechanisms^8^, locus-targeted antigen gene rearrangement occurs in many different bacteria, fungi and protists^9, 10^. Perhaps surprisingly, the processes that initiate antigen gene recombination are poorly characterised in all but *Neisseria gonorrhoeae* ^11, 12^, *Trypanosoma brucei* (see below) and, to some extent, *Borrelia burgdorferi* ^13, 14^.

Antigenic variation in *T. brucei* relies upon continuous changes in the identity of the Variant Surface Glycoprotein expressed on the parasite cell surface^15^. A single *VSG* gene is transcribed, by RNA Polymerase (Pol) I, in a parasite cell from one of around 15 telomeric loci termed bloodstream expression sites (BESs)^16, 17^. In these complex multigene transcription sites, a *VSG* appears always to be the most telomere-proximal gene and is separated from a variable number of upstream expression site associated genes (*ESAG*s)^18^ by an array of 70-bp repeats. Such repeats are also found in smaller numbers upstream of thousands of silent *VSG* genes and pseudogenes that are mainly located in arrays in the subtelomeres of 11 megabase-sized chromosomes^19–24^. *T. brucei* can activate a silent *VSG* using HR, most commonly by gene conversion^25^, where the donor *VSG* sequence is copied into the actively transcribed BES and replaces the previously expressed *VSG*. The 70-bp repeats frequently demarcate the upstream extent of *VSG* gene conversion. *VSG* switching can also occur by turning off complete transcription at the single transcribed BES and turning on complete transcription from one of the other previously silent BES. The mechanisms behind transcriptional switching remain elusive, but there is no evidence this process involves recombination. *VSG* switching is impaired when the key enzyme of HR, RAD51, is mutated^25–28^, an effect mirrored after mutation of other factors that aid homology-directed strand exchange by HR, including BRCA2 ^29, 30^ and at least one RAD51-related protein (paralogue)^31, 32^.

How HR is targeted to the active BES to cause recombinational activation of a silent *VSG* is an unresolved question^33^. Engineering the targeting of yeast I-SceI endonuclease activity within the BES in the vicinity of the *VSG* elicits *VSG* recombination^27, 34, 35^, demonstrating that a DNA double strand break (DSB) can be a trigger of *VSG* switching. However, no native BES-focused nuclease has been described, and though ligation-mediated PCR has detected breaks in the BESs, this assay has not revealed evidence for a discrete break location within, or indeed limited to, the active BES ^27, 28, 34^. Loss of telomere length or protection can also result in BES breaks and *VSG* switching ^28, 36, 37^. Finally, mutation of the RNA-DNA endonucleases RNase H1 or RNase H2 leads to VSG switching and BES damage ^38–40^, effects also seen after loss of the DNA damage signalling kinase ATR ^41^. One potential explanation to connect RNA-DNA hybrids and *VSG* switching is the demonstration that the actively transcribed BES is replicated earlier than all silent BESs ^42, 43^, but several aspects of any model that might explain the roles of RNA-DNA hybrids are unclear. One key question is how RNA-DNA hybrids contribute to *VSG* recombination switching; are they the cause of DNA breaks in the BES, or do they form in response to breaks? RNA-DNA hybrids are widespread epigenetic features of all genomes and can be resolved by RNase H-mediated degradation of the RNA within the hybrids ^44^. R-loops are a particular form of RNA-DNA hybrids in which RNA base-pairs with one strand of the DNA helix, displacing the other DNA strand. R-loops can be generated during transcription, but further activities are emerging in all genomes (Santos-Pereira and Aguilera, 2015, including initiation and arrest of replication initiation ^45, 46^, transcription activation and termination ^47, 48^, chromatin formation ^49, 50^, and telomere function ^36, 51^. In many such activities, R-loops can lead to genome instability ^52–54^, at least in part by generating DNA breaks, such as during class switch recombination in mammalian B lymphocytes ^55, 56^. However, it is also increasingly clear that RNA-DNA hybrids can form in response to DNA DSBs ^57–59^, though it is unclear in what circumstances they inhibit or promote repair of such lesions by HR.

Here, we demonstrate that *T. brucei* RAD51 interacts with RNA-DNA hybrids, and that loss of the recombinase causes genome-wide changes in R-loop abundance. Moreover, we demonstrate widespread co-localisation of R-loops with DNA DSBs, a relationship that is impaired by RAD51 loss. Finally, we demonstrate that RAD51 is needed to repair highly abundant and localised DNA breaks at the *VSG* in the single active BES, and that loss of RAD51 alters R-loop distribution both within the BES and across the silent *VSG* archive. Our data therefore reveal both generalised roles for R-loops in RAD51-directed HR in *T. brucei* and uncover specialised features of this activity in the critical immune evasion reaction of antigenic variation.

## Results

### *T. brucei* RAD51 binds RNA-DNA hybrids

Our previous work on *T. brucei* RNaseH1 and RNaseH2 revealed a connection between R-loops in the *VSG* BESs and damage-induced *VSG* switching ^38, 39^. To test and explore this connection, we performed DNA-RNA hybrid immunoprecipitation (DRIP) with S9.6 antiserum from nuclear extracts of both bloodstream form (BSF, mammalian host stage) and procyclic form (PCF, insect vector stage) *T. brucei* and identified interacting proteins by mass-spectrometry (MS) ^60, 61^. 602 proteins were recovered in at least one of six DRIP-MS replicates from the two life cycle forms (Girasol et al, BIORXIV/2023/540366); amongst these putative RNA-DNA hybrid interactors, RAD51 was detected in DRIP-MS from BSF cells but not PCF cells (Fig.1A). To test this predicted RAD51-R-loop interaction, we performed DRIP in an *RNase H1*-/- (null) mutant, where RNA-DNA hybrids could be more readily precipitated, and the recovered chromatin was separated and analysed by western blot with anti-RAD51 antiserum. RAD51 signal was more abundant in the DRIP sample than in the input, and the signal was diminished upon treatment with *E. coli* RNase HI (EcRHI; Fig.1B). To ask further if RAD51 acts upon RNA-DNA hybrids, we generated *rad51*-/- mutants (Fig.1C, S1A). To do so, we engineered MiTat1.2 bloodstream form cells, which predominantly express VSG221 (also named VSG2) from BES1 (see below) ^38, 39^, to allow modification via CRISPR-Cas9 ^62^. We then compared S9.6 immunofluorescence in the Cas9-expressing parental cell line (hereafter called wild type, WT) and *rad51-/-* cells (Fig.S1B and see below) and found that the signal was significantly reduced in the mutant (Fig.1C). In both parasite lines, the S9.6 signal was depleted by treatment with EcRHI, confirming the antiserum mainly detects RNA-DNA hybrids in these conditions. These data reveal interaction of RAD51 with RNA-DNA hybrids and indicate a role for the recombinase in homeostasis of the levels of these nucleic acid structures.

**Figure 1.**
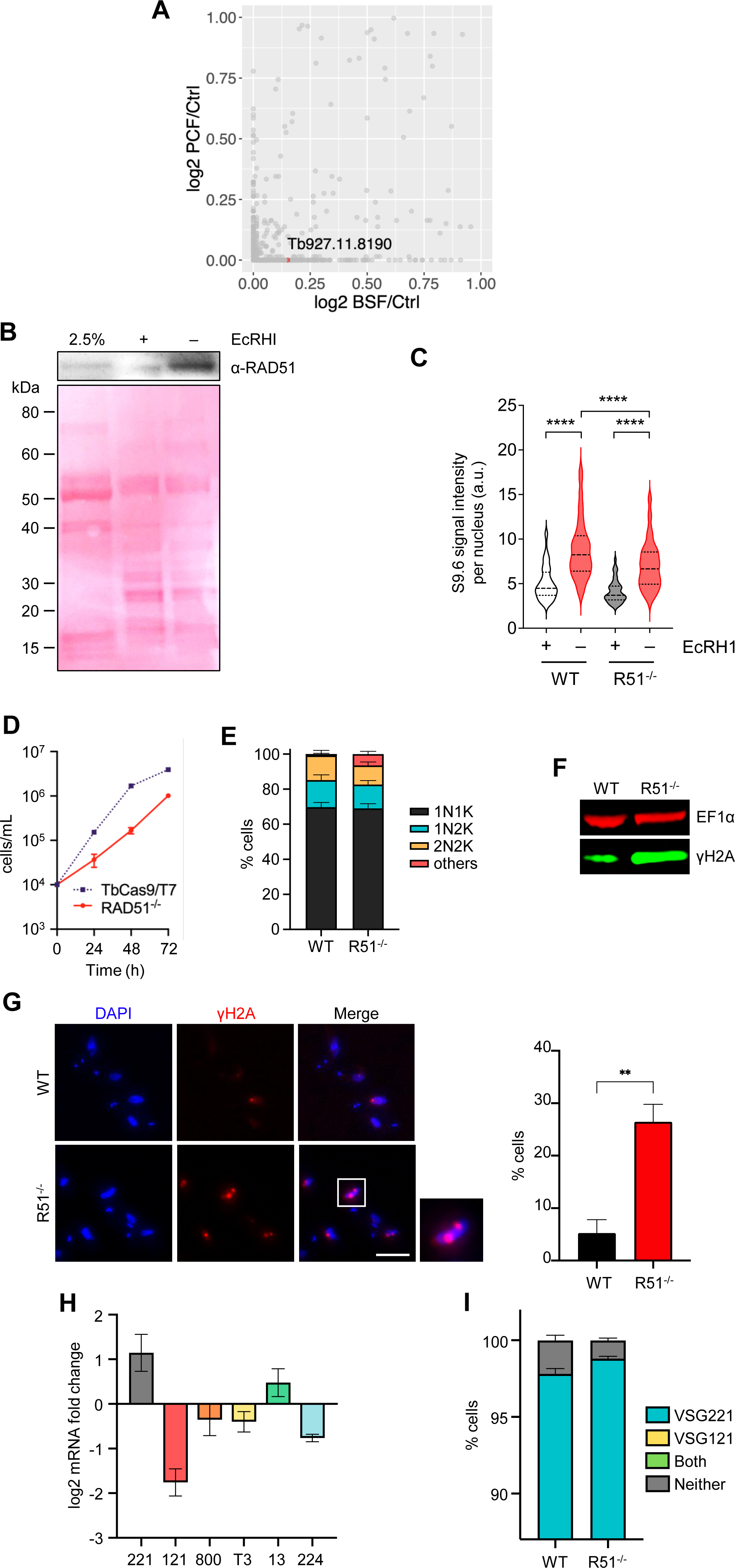
*T. brucei* RAD51, which promotes VSG switching, interacts with RNA-DNA hybrids. (A) Scatter plot of log_2_-transformed mean emPAI values of gene MS data identified from BSF and PCF S9.6 DRIPs relative to benzonase-treated controls; *RAD51* (gene ID Tb927.11.8190) is highlighted. (B) Top panel shows western blot of RAD51 in *RNaseH1-/-* cells (input) and after S9.6 DRIP, with (+) and without (-) *E. coli* RNase HI (EcRHI) treatment; bottom panel shows the same blot stained with Ponceau. (C) Violin plots show intensity of nuclear S9.6 immunofluorescence signal in WT and *rad51*-/- cells, with (+) and without (-) EcRHI treatment, in each case for > 100 cells; the median is shown by a heavily dotted line and the interquartile range by surrounding lightly dotted lines; statistical significance was determined through one-way ANOVA followed by Šídák’s multiple comparisons test: ****p < 0.0001. (D) Growth comparison of *rad51-/-* and wild type (WT, TbCas9/T7 expressing) cells; error bars represent SEM from three independent experiments. (E) Cell cycle profiles of *rad51*-/- and WT cells as determined by DAPI staining of nucleus (N) and kinetoplast (K) configurations in individual cells: 1N1K, 1N2K, 2N2K, or aberrant (others); values are shown as a proportion of >300 cells, and error bars represent SEM from three independent experiments. (F) γH2A western blot of *rad51*-/- and WT whole-cell extracts; EF1α was used as a loading control. (G) Representative microscopy images of γH2A immunofluorescence of *rad51-/-* mutants and WT cells (scale bar = 5 µm); graph shows the percent of cells with detectable γH2A signal (error bars signify SEM from three independent experiments, counting at least 50 cells in each experiment; statistical significance determined using t-test, **p < 0.01). (H) RT-qPCR quantification of different *VSG* mRNA levels in *rad51*-/- mutants compared to WT cells; error bars indicate SEM of two independent experiments. (I) Graphical representation of VSG immunofluorescence analysis showing the proportion of *rad51*-/- and WT cells expressing VSG221 alone, VSG121 alone, both VSG221 and VSG121, or neither; error bars represent SEM from three independent experiments, counting >300 cells in each experiment.

### Loss of RAD51 reduces VSG switching

To test if *rad51-/-* cells generated by CRISPR-Cas9 are comparable to previously described mutants, we tested for several phenotypes. First, the *rad51-/-* mutants displayed a growth defect relative to their WT cells (Fig.1D), consistent with previous reports ^26^. An increased proportion of cells with abnormal nucleus to kDNA (mitochondrial genome) ratios (‘others’; Fig.1E) was seen in the mutants, a perturbation that did not result from Cas9 expression. Loss of RAD51 also resulted in increased levels of nuclear DNA damage, here assessed via the detection of phosphorylated histone H2A (yH2A) ^63^ both by western blot (Fig.1F) and immunofluorescence (Fig.1G).

Next, we tested if loss of RAD51 affects VSG switching. In the Cas9 WT cell line here used, a small proportion of cells (2.18±0.34%) switched off expression of *VSG221* (transcribed from BES1) when replicating in culture (Fig.1I), as seen in other WT cells ^38, 39, 41, 64, 65^. We tested if this low level of stochastic switching was affected in the *rad51-/-* parasites: RT-qPCR revealed increased numbers of cells in the population that were transcribing *VSG221*, with a concomitant reduction in cells expressing 4 of 5 *VSG*s located in predominantly silent BESs (Fig.1H); in addition, immunofluorescence with antiserum against VSG221 and VSG121 (expressed from predominantly silent BES3) revealed fewer cells in the mutant that did not express either protein (1.19±0.15%) (Fig.1I), and hence fewer cells that had switched to a distinct, non-probed, VSG. These data confirm previous observations, using distinct assays ^26^, that deletion of *RAD51* impairs VSG switching. Notably, the effects of RAD51 loss on VSG switching are distinct from the increased VSG switching (observed using the same assays) after loss of either RNase H1 or RNase H2A ^38, 39^.

### Loss of RAD51 alters R-loop distribution in the *T. brucei* genome core

To ask if R-loop localisation is altered by loss of RAD51, DRIP coupled with deep sequencing (DRIP-seq) was performed using WT cells, *rad51*-/- mutants and *RNase H1*-/- mutants ^39, 66^[40,68], and the sequenced DNA in the recovered RNA-DNA hybrids was then mapped to the *T. brucei* genome. DRIP specificity was confirmed by mapping DRIP-seq reads from two independent experiments in *rad51-/-* mutants to the CDS of all RNA Pol II-transcribed genes, in each case with and without treatment with EcRHI (Fig.S2A). A decrease in the DRIP-seq signal within the CDS (discussed below) was seen in both replicates in EcRHI- treated versus -untreated experiments (Fig.S2A). In addition, qPCR of DRIP for selected loci confirmed EcRHI sensitivity (Fig.S2B,C).

We first examined the impact of RAD51 loss across the core genome that comprises multigenic (polycistronic) transcription units (PTUs). In previous work, we showed that R-loops are enriched at the start and ends of the PTUs, and accumulate within the PTUs at sites of pre-mRNA editing ^66^]. RAD51 loss had two clear effects associated with RNA Pol II transcription. First, the distribution of R-loops within the transcribed PTUs was altered in *rad51-/-* mutants (Fig.2A,B): in contrast to WT cells and *RNase H1-/-* mutants, where DRIP-seq signal was enriched in the flanks of the CDSs, *rad51-/-* mutants showed greater levels of DRIP-seq enrichment within the CDS (Fig.2B; a change confirmed by DRIP-qPCR: Fig.S2C). Thus, loss of RAD51 affects the localisation of R-loops at sites of trans-splicing and/or polyadenylation. Second, *rad51-/-* mutants displayed a marked accumulation of DRIP-seq signal at regions in WT cells that show enrichment for histone variant H2A.Z (Fig.2A), a marker of PTU transcription start sites (TSS)^67^. Though R-loops were previously shown to localise at PTU TSSs ^66^, the level of DRIP-seq enrichment at these loci in *rad51-/-* mutants appeared more pronounced than in WT cells or *RNase H1-/-* mutants (Fig.2A, C), an effect not seen at transcription termination sites (TTSs). Furthermore, comparing DRIP-seq pattern in the *rad51-/-* mutants relative to H2A.Z ChIP-seq in WT cells across regions 20 Kb upstream and downstream of all TSSs revealed considerable similarity (Fig.2D): a Pearson correlation coefficient of 0.9785 (p < 0.0001) was seen between the two data sets, representing a more significant correlation than seen when comparing H2A.Z signal in WT cells with DRIP-seq in either *RNase H1-*/- or WT cells (Fig.S3A). Since the DRIP-seq enrichment in the *rad51-/-* mutants was reduced by treatment with EcRHI (Fig.S3B), we conclude that *T. brucei* RAD51 plays a hitherto undetected role in the deposition and/or regulation of R-loops associated with RNA Pol-II transcription initiation.

**Figure 2.**
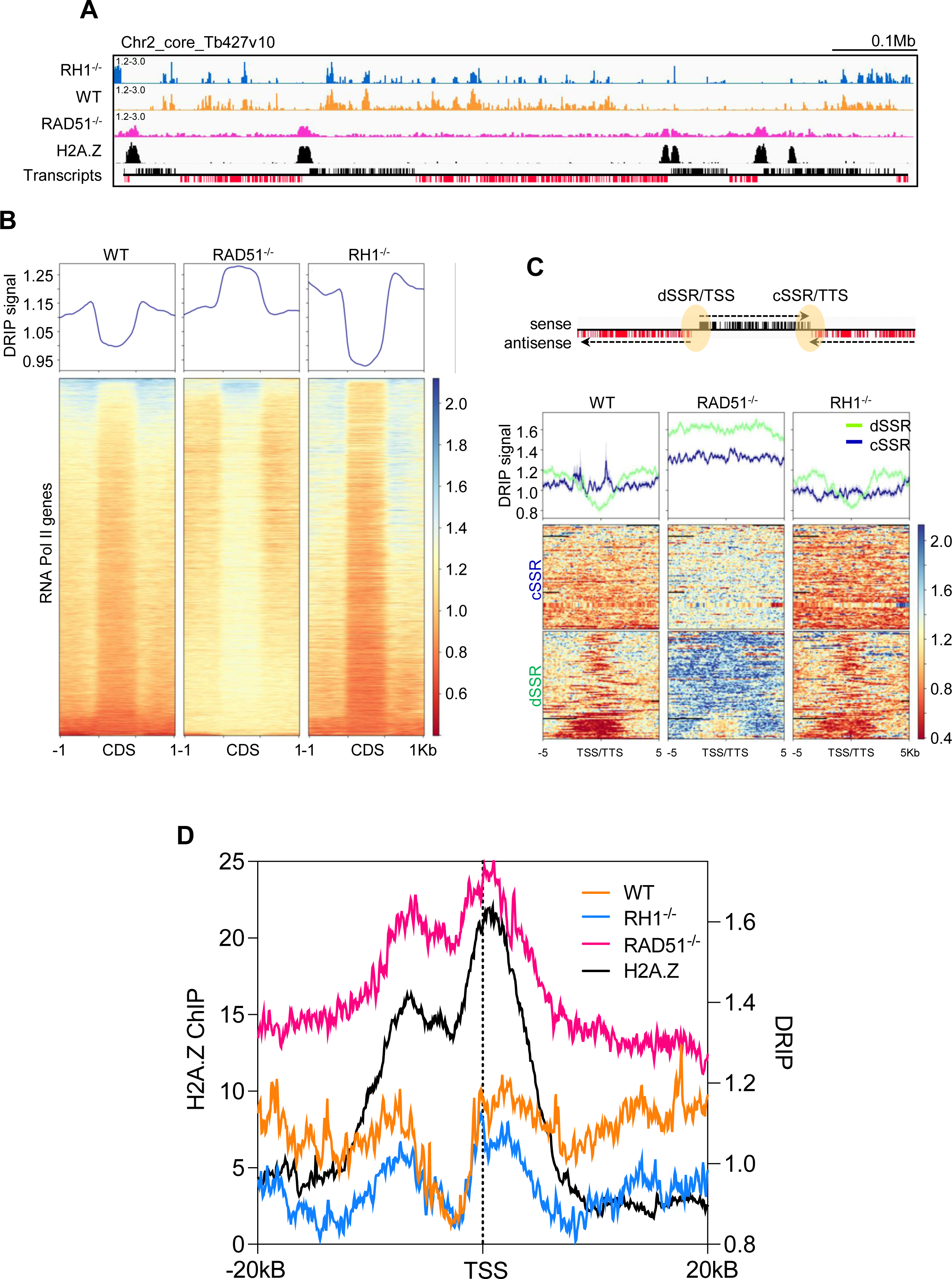
Loss of *T. brucei* RAD51 alters R-loop distribution within and at the transcription start sites of RNA Polymerase II multigene transcription units. (A) DRIP-seq read mapping (DRIP/input) across a selected region of chromosome 2 in *RNase H1-/-* (*RH1*-/-, blue), wild-type (WT, orange) and *rad51*-/- (magenta) cells, compared with H2A.Z ChIP-seq in WT cells (black); the bottom track shows genes with sense (black) and antisense (red) transcripts. (B) Upper panel: metaplot of DRIP-seq signal (relative to input) across all RNA Pol II–transcribed genes (scaled to the same size), including the 1 kb sequences that flank their CDS. Lower panel: heatmap of DRIP-seq signal fold change across all CDS +/- 1 kb. (C) Metaplots (top panel) and heatmaps (bottom panel) of WT, *rad51*-/- and *RH1*-/- DRIP-seq signals across convergent (blue) and divergent (green) strand switch regions (cSSR and dSSR, respectively) between all RNA Pol II PTUs; the loci are aligned on their centres and include 5 kb of upstream and downstream sequence; the cartoon illustrates that cSSRs are transcription termination sites (TTSs), and dSSRs are transcription start sites (TSSs). (D) Metaplots of H2A.Z ChIP-seq signal (black, left y-axis) in WT cells is compared to DRIP-seq signal (right y-axis) in WT (orange), *RH1*-/- (blue) and *rad51*-/- (magenta) (right y-axis) cells over 40 kb regions surrounding all predicted TSSs. All metaplots show mean plus SEM (shaded).

Pronounced enrichment of R-loops in the *T. brucei* genome is also seen at centromeres [68], which are found in the transcribed core of chromosomes 1-8 ^68^ and in the non-transcribed subtelomeres of chromosomes 9-11 ^69^. To date, the size and detailed sequence of the centromeres is uncertain ^69, 70^. To clarify centromeric enrichment of R-loops, we used long-read Nanopore sequencing to re-sequence the Lister 427 genome. 23 Nanopore contigs were identified that encompass at least part of the centromeres found in each chromosome (Fig.3), and DRIP-seq mapping to these contigs confirmed widespread localisation of R-loops in both WT cells and in the *RNase H1-/-* mutant. In the *RNase H1* -/- cells DRIP-seq signal was more pronounced in around 60% of the centromeres, for reasons that are unclear (Fig.3B). In the *rad51-/-* mutants DRIP-seq signal was less pronounced in the centromeres that showed most enrichment in the *RNase H1*-/- mutants (Fig.3A, B), indicating that here too the recombinase influences R-loop localisation.

**Figure 3.**
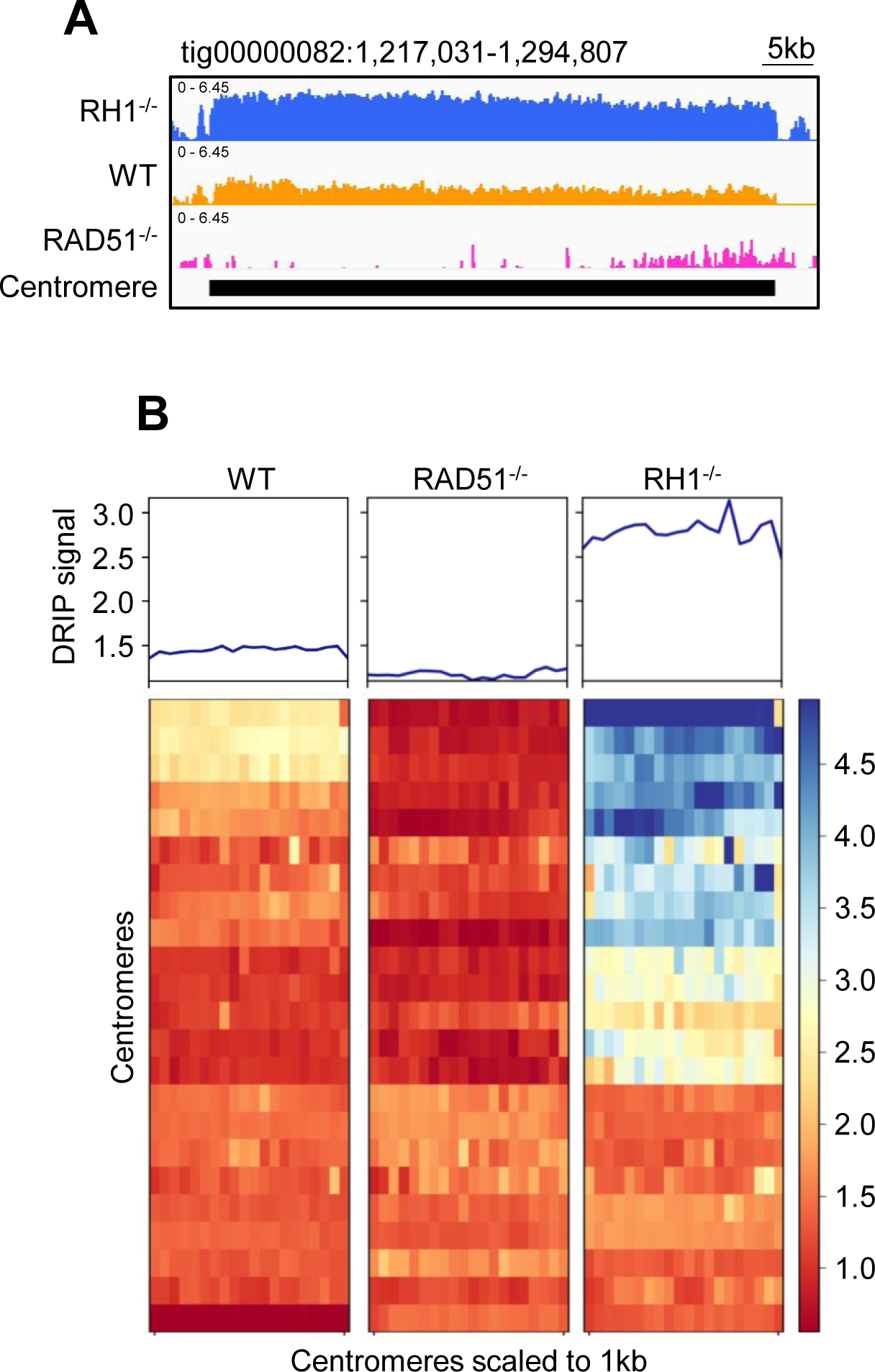
Loss of *T. brucei* RAD51 alters R-loop localisation to centromeres. (A) DRIP-seq read mapping (DRIP/input) across one Nanopore contig covering a complete centromere (black box), comparing *RNase H1-/-* (*rh1-/-*, blue), wild-type (WT, orange) and *rad51*-/- (magenta) cells. (B) Metaplots (top panel) and heatmaps (bottom panel) of wild type, *rad51*-/- and *rh1*-/- DRIP-seq signals (relative to input) across all centromere-containing contigs (scaled to the same size); data shows mean plus SEM (shaded).

### Loss of RAD51 alters R-loop distribution in the *VSG* expression sites

We next examined DRIP-seq mapping in the *VSG* BESs (Fig.4). Modest DRIP-seq enrichment was seen in silent or active BESs in WT cells, with somewhat greater signal detected downstream of the *VSG,* while DRIP-seq enrichment was greater across the active and silent BESs in *RNase H1*-/- mutants (Fig.4A, B)^39^. Loss of RAD51 had two effects on DRIP-seq distribution. First, and most pronounced, was reduced enrichment towards the telomere of the BESs (Fig.4B) in *rad51-/-* cells, a change accounted for by reduced DRIP-seq signal across the 70-bp repeats, where DRIP-seq enrichment strikingly increased in the *RNase H1*-/- mutants (Fig.4C)^39^ relative to WT. Second, somewhat increased DRIP-seq enrichment was seen across the *ESAG*s upstream of the 70-bp repeats, though this change was not as extensive as seen in *RNase H1*-/- cells (Fig.4D, Fig.S2B). When comparing DRIP-seq mapping across the *VSG* genes in the BESs (Fig.4E) or the telomeric repeats (Fig.4F), any effects of RAD51 loss were less apparent than after RNase H1 loss, though R-loops were detectable in the *VSG*s (Fig.S2B). Taken together, these data indicate that RAD51 is predominantly involved in the formation or stabilisation of R-loops at *VSG*-associated 70-bp repeats but not at telomeres, unlike TERRA in other eukaryotes ^71, 72^.

**Figure 4.**
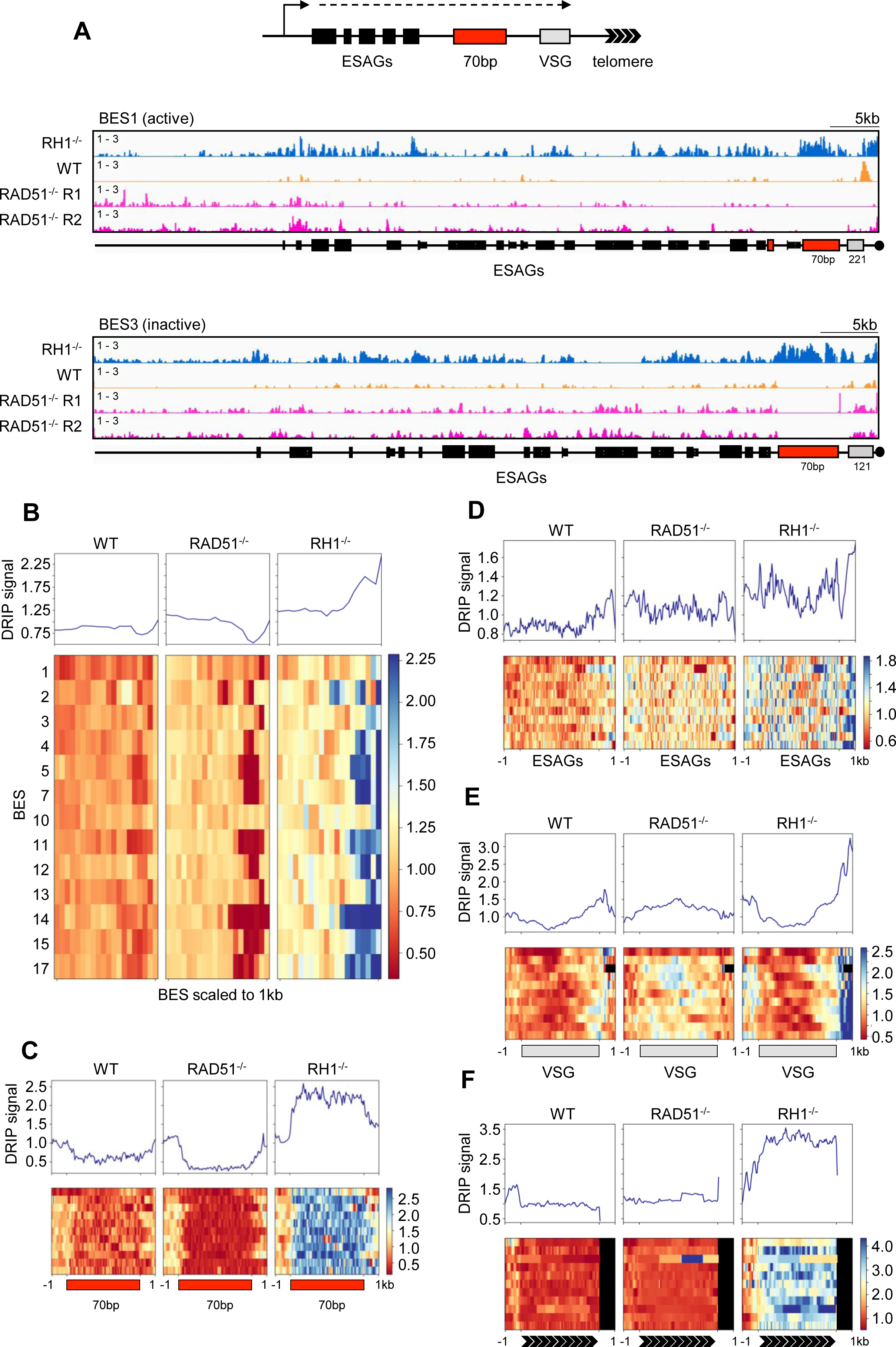
Loss of *T. brucei* RAD51 alters R-loop localisation in VSG expression sites. (A) A schematic representation of a bloodstream expression site (BES) is shown and, below, DRIP-seq read mapping (DRIP/input) across BES1 and BES3, comparing *RNase H1-/-* (*rh1-/-*, blue), wild-type (WT, orange) and two independent *rad51*-/- (magenta) cells. Metaplots (top panel) and heatmaps (bottom panel) of WT, *rad51*-/- and *rh1*-/- DRIP-seq signals (relative to input) are shown for all BESs (B), or focusing on the 70-bp repeats (C), *ESAG*s (D), *VSG*s (E) and telomeres (F); in all cases, the BESs or BES features are scaled to the same size and data for silent BESs shows mean plus SEM (shaded).

### Loss of RAD51 or RNase H1 alters R-loop distribution at silent, subtelomeric array *VSG*s

To ask if RAD51’s influence on *VSG*-associated R-loops is limited to the *VSG* BESs, we next examined the subtelomeric *VSG* arrays, again using Nanopore long-read contigs, which allowed us to clearly assess R-loop localisation in this component of the genome for the first time (Fig.5). In WT cells, a similar distribution of DRIP-seq mapping to that observed around RNA Pol-II transcribed CDSs (Fig.2B) was seen in the array *VSG*s (Fig.5A,B), and this pattern became exaggerated in *RNase H1-/-* mutants (Fig.5A,B). Thus, R-loops accumulate around potentially all subtelomeric *VSG* genes, irrespective of their transcription status. Loss of RAD51 altered the R-loop distribution, with *rad51-/-* mutants showing greater DRIP-seq enrichment within the CDS than in the flanks (Fig.5A,B), an effect mirroring what was seen in RNA Pol-II transcribed genes in the genome core (Fig.2A,B). Together, these results indicate that *VSG*-associated R-loops, whose formation and/or stability is modulated by RAD51 and RNase H1, are not only seen in telomere-adjacent *VSG*s within transcription sites but are also widespread across the extensive and mainly silent subtelomeric *VSG* archive.

**Figure 5.**
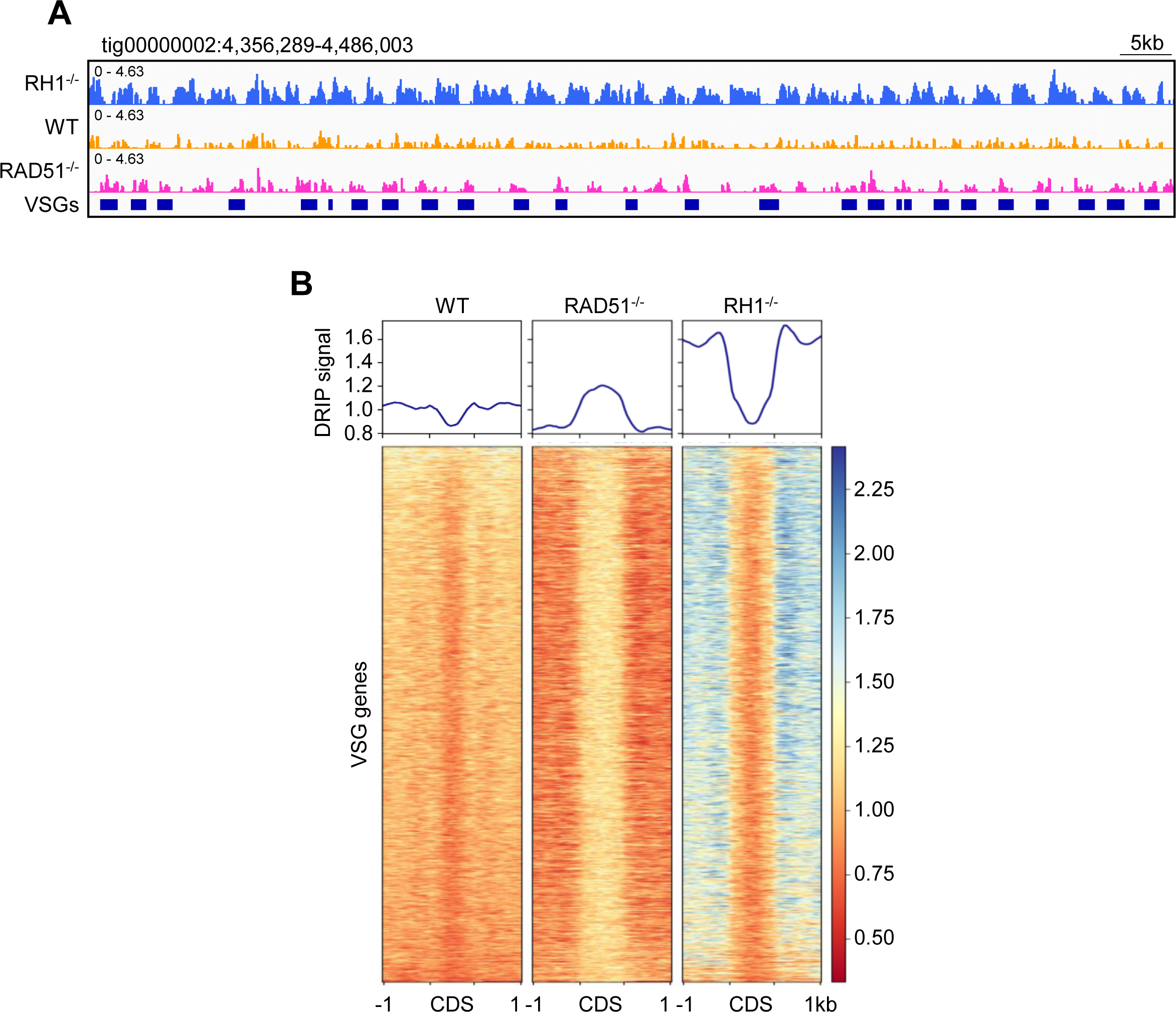
Loss of *T. brucei* RAD51 alters R-loop localisation around transcriptionally silent subtelomeric array *VSG*s. (A) DRIP-seq read mapping (DRIP/input) across one Nanopore contig containing an array of subtelomeric *VSG*s, comparing *RNase H1-/-* (*rh1-/-*, blue), wild-type (WT, orange) and *rad51*-/- (magenta) cells. (B) Metaplots (top panel) and heatmaps (bottom panel) of wild type, *rad51*-/- and *rh1*-/- DRIP-seq signals (relative to input; mean is shown and SEM is shaded) across the predicted CDSs of all identified subtelomeric *VSG*s (scaled to the same size), plus 1 kb of upstream and downstream sequences.

### DNA breaks mapped by BLISS reveal association with RAD51-directed R-loops

Given the above data reveals a link between R-loops and RAD51, a key enzyme that directs repair of DNA damage, we next sought to map the locations of DNA breaks in the *T. brucei* genome. To do so, we used breaks labelling in situ and sequencing (BLISS), where DSBs in a cell population are mapped across the genome via adapter ligation, *in vitro* transcription and RT-PCR amplification ^73^. To ask which breaks are acted upon by RAD51, we compared BLISS mapping in the WT cells and in the *rad51*-/- mutants derived from them.

A total of 17,134 and 11,160 BLISS ‘peaks’ were predicted using MACS2 ^74^] in the WT and *rad51*-/- populations, respectively, with the distribution across a range of genomic landmarks being very comparable (Fig.S4A). It should be noted that these BLISS peaks represent sites of read accumulation on one DNA strand, meaning they do not necessarily represent discrete two-ended DNA DSBs (which would have flanking reads on both the Watson and Crick strands) and may instead be single ends of highly processed DSBs or one-ended DSBs ^73, 75^. 69% and 71% of BLISS reads were derived from the rRNA loci in WT and *rad51*-/- cells, respectively, suggesting these are pronounced sites of DNA breaks, consistent with other organisms ^76, 77^. As the mapping removes reads of low quality, including from highly repeated sequences such as the rRNA, this abundance is not reflected in the genome-wide distribution plots (Fig.4A) but, nevertheless, the small remaining rRNA BLISS signal was more commonly found here after loss of the recombinase (0.13% in WT, 0.25% in *rad51-/-*; Fig.S4A), suggesting at least some rRNA gene breaks are acted upon by RAD51-directed HR. To explore how the BLISS peaks are distributed elsewhere, metaplots of BLISS reads were generated for other genomic features. Centromeres and tRNAs were flanked by BLISS peaks in both WT cells and *rad51*-/- mutants (Fig.S4B). Most BLISS peaks were detected within the PTUs (‘others’, Fig.S4A) and metaplots of the genes revealed a somewhat diffuse arrangement of increased BLISS signal flanking the RNA Pol-II–transcribed CDSs (Fig.S4B). As this signal appeared less abundant in *rad51-/-* mutants, such mapping may be a signature of lesions repair by means other than HR, such as transcription-coupled nucleotide excision repair ^78^. In contrast to other organisms, where DSBs are strongly associated with TSSs ^79^, we saw no clear BLISS signal localization at the start of the PTUs but did detect signal at TTSs in both WT and *rad51*-/- cells (Fig.S4C). The three strongest BLISS peaks (Fig.S4D) mapped to unitig851_maxicircle_Tb427v10, which represents mitochondrial maxicircles that are known to be found frequently as linear molecules after the generation of DNA DSBs due to topoisomerase II ^80^, and hence provide an example of correspondence between BLISS signal and known DNA breaks.

To ask if R-loops correlate with DSBs across the *T. brucei* genome, metaplots were generated for all loci where BLISS peaks were identified by MACS and were compared with DRIP-seq distribution at the same loci in both WT and *rad51-/-* cells (Fig.6). DRIP-seq signal showed a pronounced enrichment that centred on the more discrete BLISS signal in WT cells, indicating break sites frequently associate with RNA-DNA hybrids. On the other hand, in *rad51*-/- mutants DRIP-seq enrichment was depleted at the centre of the BLISS signals. These data suggest that loss of RAD51 impairs R-loop localisation at BLISS-mapped breaks, indicating the recombinase acts to recruit or stabilize the hybrids at many DNA break sites, potentially during the process of DSB repair.

**Figure 6.**
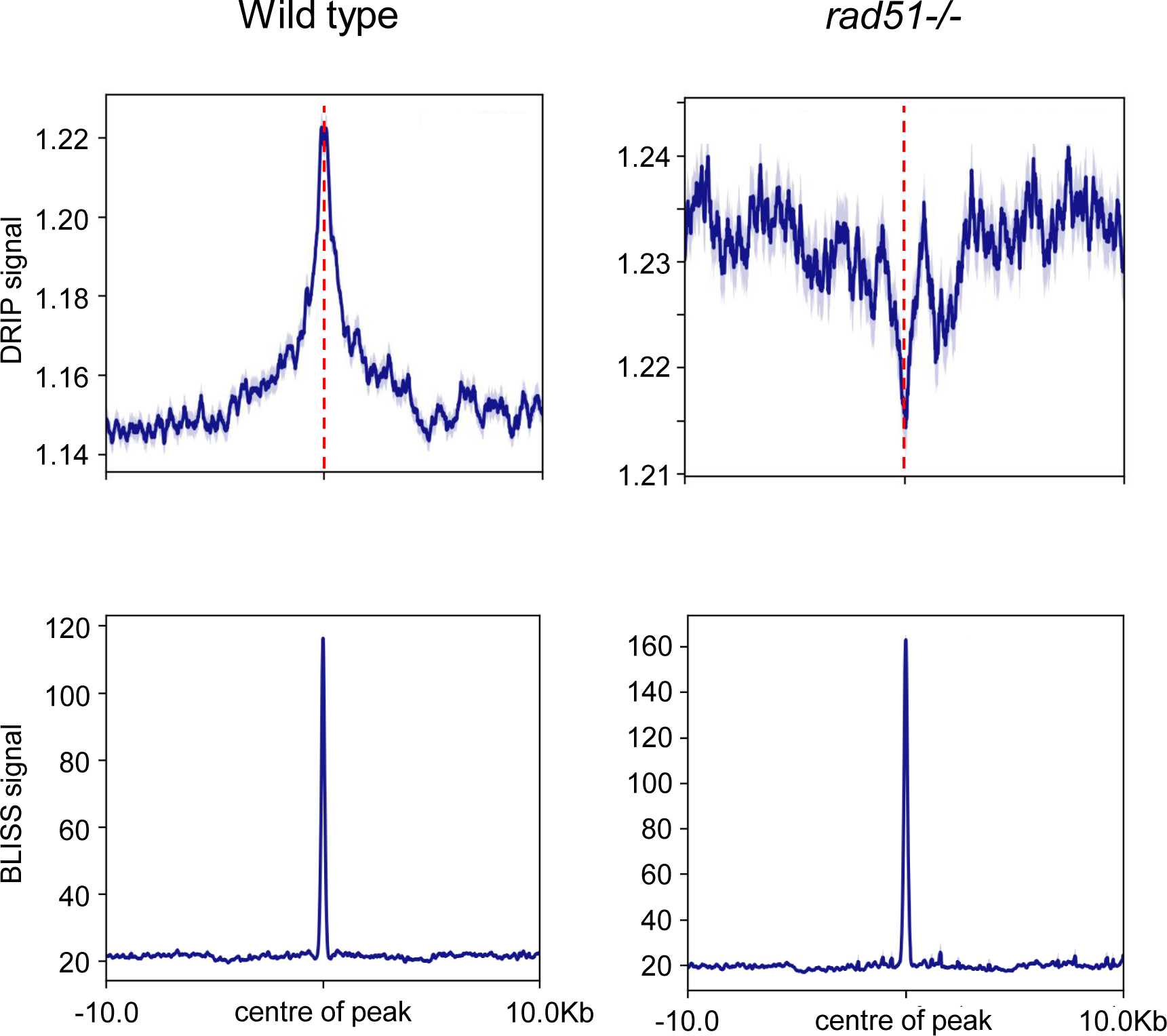
R-loop accumulation at DNA breaks is perturbed by loss of RAD51 across the *T. brucei* genome. Metaplots showing (upper panels) the distribution of DRIP signal (DRIP/input) relative to (lower panels) BLISS coverage (BLISS signal is read count per million mapped reads) in wild type and *rad51*-/- cells. DRIP and BLISS signals are the mean plus SEM (shaded) aligned on the centre of the BLISS signal (dotted line) and including 10 kb of upstream and downstream sequence is shown.

### The transcribed *VSG* expression site contains a *VSG*-localised DNA break acted upon by RAD51

To ask if breaks can be detected in the *VSG* BESs, and if RAD51 loss impacts on the level and distribution of such breaks, BLISS reads were mapped to all annotated BESs (Fig.7A). This mapping revealed three findings. First, BLISS signal was strikingly higher in active BES1 than all silent BESs. Second, BLISS accumulation was most pronounced proximal to the telomere, and particularly at a relatively discrete location around the 3’ end of the *VSG221* gene. Third, the incidence of this most pronounced *VSG221-*associated BLISS signal increased in the *rad51*-/- mutants compared with WT. Quantification of the BLISS signals illustrates these findings. The most prominent, *VSG221*-localised BLISS peak in BES1 had ∼30x higher BLISS signal in WT cells and ∼197x higher in the *rad51-/-* mutant when compared with the average BLISS signal from all inactive BESs at the same location (Fig.7A). In addition, whereas this main BES1 BLISS peak was ranked 119^th^ in terms of BLISS signal in WT cells, it was ranked as 4^th^ highest in the *rad51*-/- mutant (Fig.S4D, surpassed only by three mitochondrial maxicircle peaks). These data indicate the presence of a putative DNA DSB or multiple DSBs in the telomere-proximal end of the active *VSG* BES, suggesting an association with active transcription and that the repair of such breaks requires RAD51. Furthermore, while the BLISS mapping indicates putative DSBs located within the active BES that are consistent with previous suggestions of telomere-proximal BES regions being ‘fragile’ ^27^, one 3’ *VSG*-focused region in the active BES is the location of most BLISS signal accumulation.

**Figure 7.**
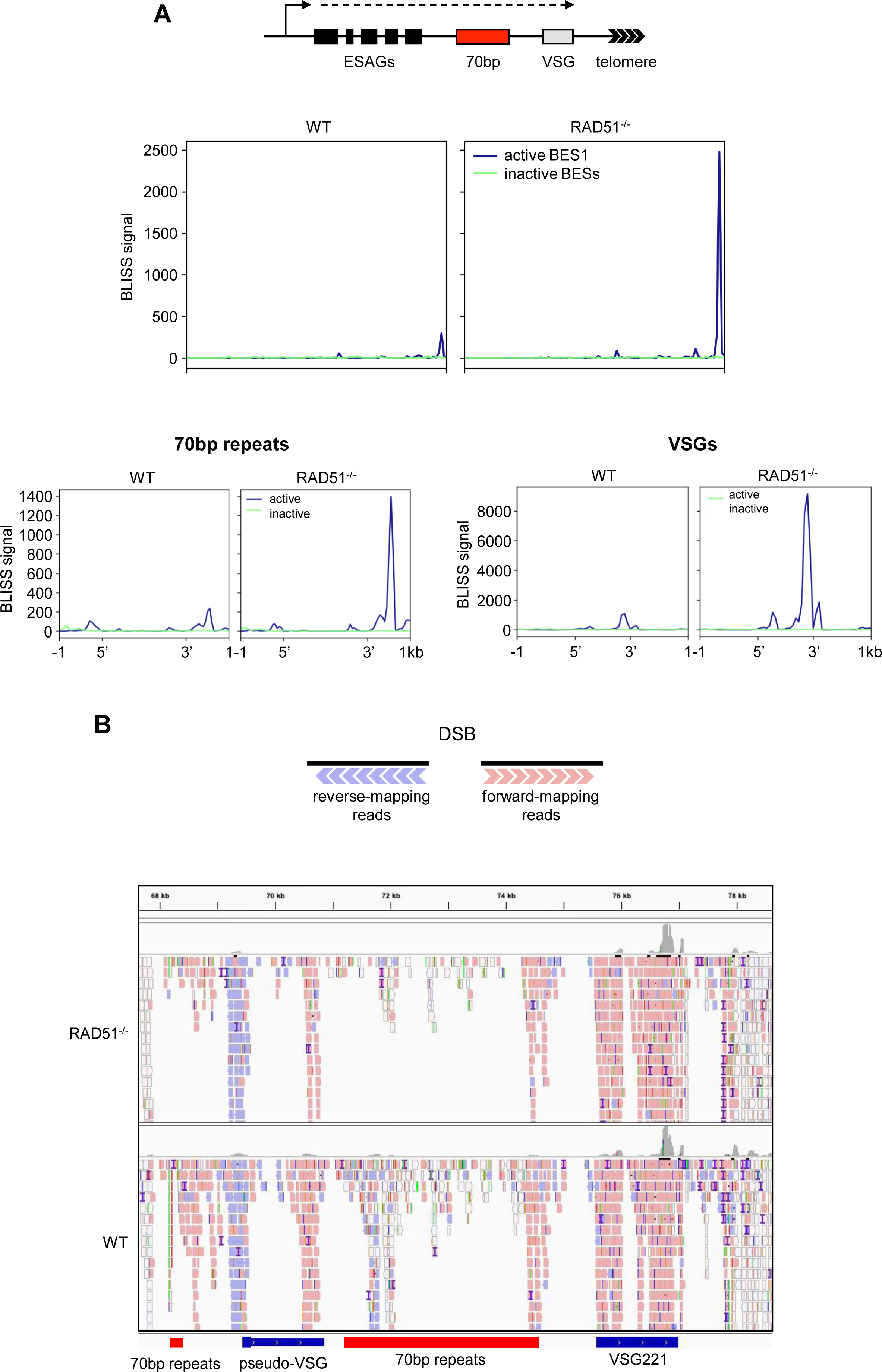
Pronounced DNA breaks at the 3’ end of the actively expressed *VSG* are repaired by RAD51. (A) Metaplots comparing the distribution of BLISS reads mapped to the active (blue, BES1) and all inactive (green) *VSG* expression sites (BESs; mean, plus SEM shaded) in wild type (WT) and *rad51-/-* cells; BLISS signal is read count per million mapped reads (data are shown as averaged, normalised signal across the features of interest). A schematic of a BES is shown and BLISS mapping across the whole length of the BESs is shown in the main panel, while the lower panels show BLISS signal across the 70-bp repeats (left) and BES-housed *VSG*s (right); in all cases the loci are scaled to be the same size. (B) The schematic shows the predicted orientation of mapped BLISS reads around a discrete DSB, and the figure below details the distribution of forward- and reverse-mapping reads at the telomere-proximal region of active BES1 in both WT and *rad51*-/- cells; locations of *VSG221*, a *VSG* pseudogene and 70-bp repeats are indicated.

To understand if the BLISS read accumulation in BES1 at the 3’ end of *VSG221* represents a discrete DSB, the arrangement of reads mapped to the telomere-proximal region of BES1 in WT and *rad51-/-* cells was examined (Fig.7B). In BLISS, DSBs are processed to generate blunt ends for adapter ligation, and a minimally processed, discrete DSB break results in forward- and reverse-mapping reads representing downstream and upstream flanks of the break, respectively (Fig.7B)^81, 82^. Here, BLISS mapping was not consistent with a discrete DSB at the 3’ end of *VSG221*, since the pronounced BLISS signal in this location was derived almost exclusively from forward-mapping reads, suggesting they represent the downstream end of a processed DSB. Equivalent levels of reverse mapping reads, representing an upstream end of a DSB, were not detectable in the BES in either WT or *rad51-/-* cells, other than a smaller BLISS peak between a short array of 70-bp repeats and a *VSG* pseudogene. In addition, and consistent with the BLISS metaplots (Fig.7A), there was no clear evidence of BLISS reads mapping within either the larger or smaller stretches of 70-bp repeats in BES1.

Many *VSG* genes contain a conserved 14-mer sequence within their 3’ UTR ^20^. Though the predominant BLISS signal was upstream of the *VSG221* 14-mer in BES1, enrichment centred on the 14-mer was apparent and became more abundant after loss of RAD51 (Fig.S5A). Since we saw R-loop accumulation across the *VSG* archive, and loss of RAD51 or RNase H1 altered this distribution, we asked if the 14-mer might represent a sequence-specific target for break generation common to most *VSG*s, whether active or silent. Fig.S5B shows metaplots of BLISS and DRIP-MS reads mapped around the 14-mer of all silent *VSG*s. DRIP-seq reads displayed modest accumulation around the 14-mer in WT cells, which became elevated in *RNase H1-/-* mutants and depleted in *rad51-/-* mutants, consistent with localized accumulation of R-loops. BLISS reads also displayed accumulation that correlated with the DRIP signal in WT cells, perhaps indicating sites of break formation, but the BLISS signal was not increased in *rad51-/-* mutants, suggesting any such breaks are not acted upon by the recombinase across the silent *VSG* archive. Hence, pronounced DNA break accumulation after loss of RAD51 is limited to the single actively transcribed *VSG*.

### Discussion

Immune evasion by antigenic variation in *T. brucei* is driven by locus-specific recombination of *VSG* genes. Available evidence indicates RAD51-directed HR plays a major role in such *VSG* gene switching, but how the reaction is targeted and triggered has been the subject of debate ^33^. Here we reveal that RAD51 plays a role in repairing DNA breaks localised to the single transcribed *VSG*, and that such a role involves the recruitment of RNA-DNA hybrids. Our data suggest that such a dedicated role in VSG switching is one specialised example of *T. brucei* RAD51 functioning genome-wide to localise RNA-DNA hybrids to DNA breaks, consistent with growing evidence of R-loop activities during DNA break repair ^57–59^.

Rad51 from mammals and yeast has been shown to bind RNA-DNA hybrids *in vivo* and *in vitro* ^72, 83^. Here, DRIP-MS and DRIP-western blot data indicate *T. brucei* RAD51 also interacts with R-loops, but whether this interaction is direct or indirect is unclear. The reduction of S9.6 nuclear signal in *rad51-/-* mutants indicates that loss of the recombinase has widespread effects on R-loop homeostasis. Such a suggestion is consistent with our analysis showing a global association between BLISS-predicted DNA breaks and sites of R-loop accumulation, and that such R-loop accumulation is impaired after loss of RAD51. Hence, this work suggests *T. brucei* RAD51 promotes R-loop localisation to DNA breaks at many loci, with the implication that RNA-DNA hybrids have so-far unexplored roles in *T. brucei* RAD51-directed DNA break repair. A single R-loop– associated repair activity across all breaks found in the parasite genome is unlikely: whereas some repair-associated R-loops in other organisms are derived from endogenous transcription ^84–86^, others have been proposed to form through non-coding RNA derived from DNA DSBs ^87–89^, and other studies have suggested *trans*-formation of R-loops ^72, 84^]. Indeed, in other organisms, such R-loop association with DNA break repair extends beyond Rad51-directed HR. Here, the pronounced association of *T. brucei* RAD51 with DSBs and R-loops we see may result from the apparent lack of the non-homologous end-joining repair pathway or RAD52 in any kinetoplastid, increasing the focus of break repair on RAD51-HR ^90, 91^.

The clearest and strongest association we detect between RNA-DNA hybrids and DNA breaks during RAD51-directed repair in *T. brucei* is in the actively transcribed *VSG* expression site (BES1), where BLISS mapping predicts the most abundant DNA break anywhere in the nuclear genome in *rad51*-/- mutants. These new data allow us to develop an emerging model for how R-loops act during *VSG* switching (Fig.8). Previously, we postulated that R-loops may be the trigger for DNA breaks that lead to *VSG* switching ^38, 39^, a role consistent with widespread evidence for the hybrids being causes of genome instability ^57, 59^. Here, we show that loss of RAD51 has two effects: reduced levels of DRIP-seq signal across the 70-bp repeats of active and inactive BESs, and increased levels of *VSG*-localised BLISS signal (indicating accumulated and/or unrepaired DSBs) in the active BES, but not inactive BESs. Hence, rather than R-loops leading to DNA damage, these data suggest that DNA breaks arise in the active BES and lead to RAD51-dependent accumulation of R-loops. In this scenario, it may be that RAD51 generates a *trans* R-loop during the break repair process and RNase H enzymes aid in their resolution, explaining the change in R-loop levels and distribution we see in the silent BESs and in the silent *VSG* arrays after loss of the RAD51, RNase H1 or H2 ^38, 39^. However, uncertainty about how and where breaks arise in the BES – discussed below – means the 70- bp repeats could be both the cause of damage and a participant in repair. Nonetheless, this model is consistent with data in other organisms suggesting transcribed loci are a focus for DNA repair, including HR ^36, 84, 92–94^, and with emerging understanding of how R-loops contribute to DSB repair.

**Figure 8.**
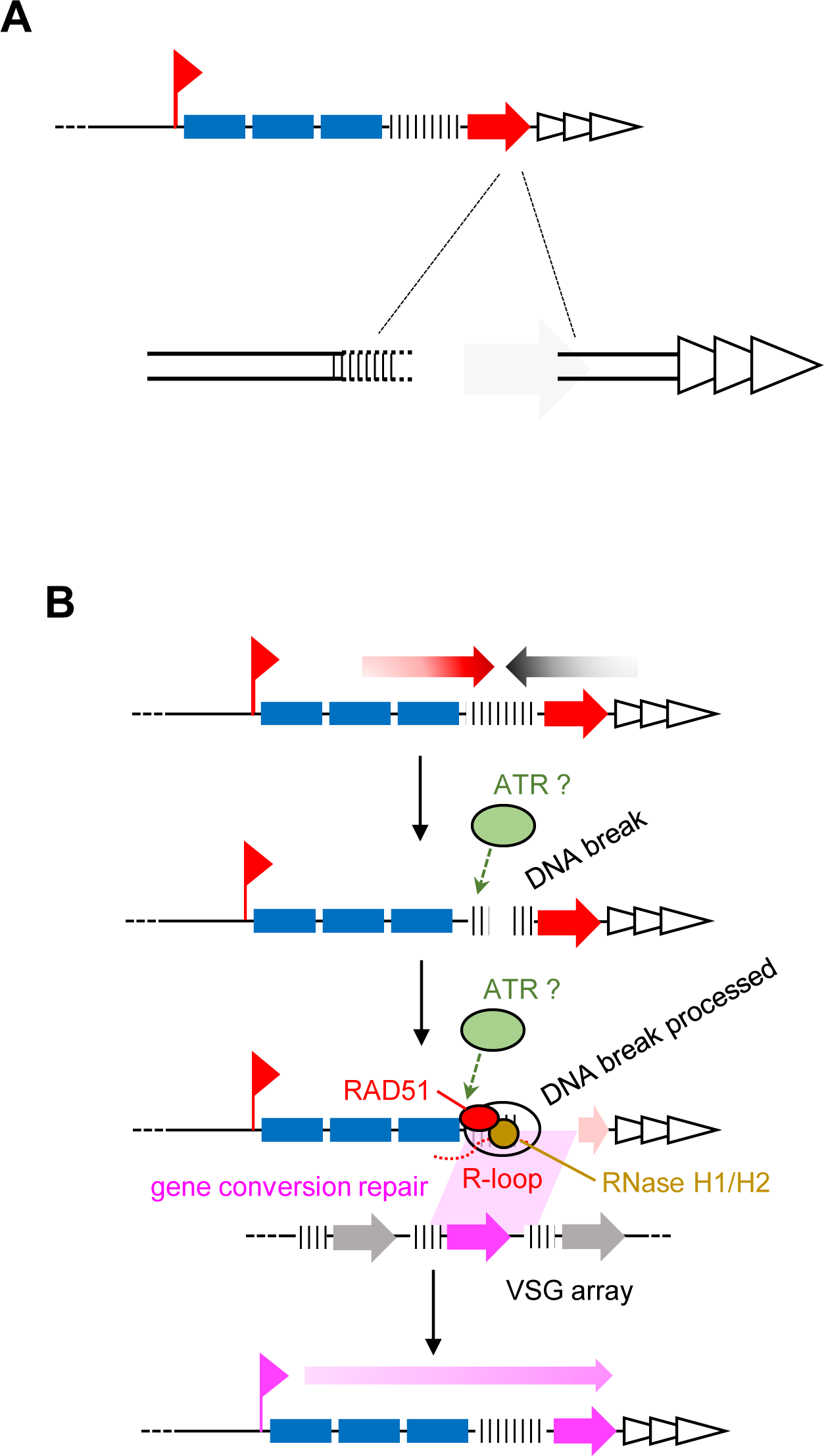
A model for the interplay of R-loops and DNA breakage during *T. brucei* VSG switching by recombination. (A) A schematic showing the BLISS-predicted location of a DNA DSB in the active *VSG* within the larger expression site. The downstream end of the break is relatively discrete, while the upstream end is more variable (dotted lines) and is predicted to reside in the 70-bp repeats. (B) A predicted order of events during VSG recombination: A DSB forms within the BES; the DSB is processed and RAD51 is recruited, directing the localisation of R-loops to the break; RAD51 (and perhaps R-loops) catalyses DSB repair via gene conversion using a silent *VSG* as template. RNase H1 and RNase H2 resolve R-loops once they form. ATR kinase recognises the DNA break and/or the R-loops.

Several aspects of this newly-refined model remain uncertain and are potentially trypanosome-specific. First, what is the nature of the DNA break(s) in the active BES, and where do they form? Though BLISS mapping reveals predominant signal accumulation around the 3’ end of the *VSG* in the active BES, the data do not support the idea that the signal here represents a discrete DSB. Instead, the predominant signal appears to correspond with the downstream end of a DNA break, with no such discrete upstream end (Fig.8A). One explanation for these data, consistent with loss of R-loops in the 70-bp repeats in *rad51-*/- mutants, is that breaks originate in the 70-bp repeats and are processed extensively, but such processing terminates at a more clearly defined region downstream of the *VSG* than upstream in the BES. It is also possible that breaks form at the 3’ end of the *VSG* and processing occurs more extensively upstream than downstream. In both cases, the downstream limitation of break processing may be the telomeric shelterin complex. Alternatively, since BLISS maps breaks in a population, the data could be explained by variation in DSB localisation with the active BES in different cells, with greater variability in their upstream location and less downstream. A complication in any such scenario is that BLISS mapping involves DSB processing, and so we cannot yet say if such putative diffuse breaks arise during BLISS sample preparation ^73^, or if extensive DSB end-processing occurs naturally in preparation for recombinational repair during *VSG* switching (Fig.8B). For similar reasons, we cannot be sure if BLISS has mapped a DSB or a distinct form of lesion, such as a stalled replication fork (Fig.8B), which is converted to a DSB during BLISS. Irrespective, the nature of the BLISS mapping appears consistent with previous attempts to detect DSBs in the BES by ligation-mediated PCR, which also appeared not to reveal a discrete break location ^27, 28, 34, 36^.

Questions also arise from this model regarding the dynamics and potential role of the R-loops in VSG switching: why does loss of either characterised nuclear RNase H enzyme increase VSG switching ^38, 39^[39,40], and are the hybrids active participants in RAD51 break repair? Increased DRIP-seq signal in *RNase H1-/-* mutants is most pronounced in the 70-bp repeats but is also seen across the BES. Furthermore, increased levels of R-loops are not limited to the active BES but is also seen in silent BESs and in silent array *VSG*s. Similarly, although elevated levels of BLISS signal in *rad51-/-* mutants is limited to the active BES, DRIP-seq changes are seen in both the active and silent BESs (and manifest as decreased levels within the 70-bp repeats but some increased signal upstream of the 70 bp repeats), and changes in R-loop distribution are seen in silent array *VSGs*. It seems likely that increased R-loop signal in the active BES after RNase H1 loss is consistent with its well-documented roles in removing hybrids [45], including at a DSB at an actively transcribed locus [97,98], in this case the BES transcribed by RNA Pol-I. What is less clear is why elevated levels of R-loops at a BES break in RNase H1 mutants might increase *VSG* switching ^39^, since some evidence suggests increased R-loop levels impede HR ^95^. However, other studies have suggested R-loop levels may alter DSB end-processing and Rad51 loading ^96–102^, which may affect repair pathway choice ^103^. In *T. brucei*, how DNA breaks that lead to VSG switching are processed is especially unclear, since mutation of the MRE11-RAD50-NBS1/XRS2 complex does not appear to alter VSG switch rate ^104^, but clearly changes *VSG* donor selection after switch induction by a targeted DSB ^64^. Impairing BES break repair by RAD51-HR may also open other routes to *VSG* switching, such as microhomology-mediated end joining ^90, 105, 106^. Increased levels of R-loops in the BES *ESAG*s after RAD51 loss may stem from more a persistent DSB(s) at the *VSG*, resulting in increased stalling of upstream transcription ^107–109^. Such stalling would increase R-loop levels and may cascade to increased transcriptional switching, resulting in R-loop formation in the previously silent BESs [40]. However, changes in DRIP-seq levels in *rad51*-/- and *RNase H1*-/- mutants in the 70-bp repeats of the silent BES may suggest a more active role for R-loops in HR during *VSG* switching. This suggestion is further supported by changes in DRIP-seq signal in the silent subtelomeric arrays: even if the silent BESs might become more transcriptionally active after mutation of RAD51 or RNase H1 (as postulated above), transcription across the very large subtelomeres is less easy to explain. In other eukaryotes several studies implicate R-loops in promoting break repair ^89, 110–112^, including evidence for the RNA providing an active role in strand exchange ^84^. It is therefore possible that the changing patterns of DRIP-seq signal we see in *rad51-/-* and *RNase H1-/-* mutants relative to WT in the silent BES and array *VSG*s reflect, respectively, decreased and increased formation of R-loops in these loci as a result of impaired or enhanced RAD51-directed homology search and strand exchange during *VSG* recombination. Thus, RAD51-directed *trans* R-loops may be widespread in *T. brucei*. However, another function for the subtelomeric RNA-DNA hybrids may yet emerge.

Unlike the actively transcribed BES, the function and formation of R-loops at centromeres is unclear, including why they might be acted upon by RAD51. Though siRNA has been found at some *T. brucei* centromeres, it has not been detected at others ^113^, perhaps explaining the uneven distribution of DRIP-seq signal we see. Equally, centromeres are not clear sites of BLISS-mapped DNA breaks, so why DRIP-seq mapping is altered in *rad51-/-* cells is unclear. Nonetheless, the damage signalling kinase ATR acts on R-loops in centromeres ^114^, and centromere-specific histones ^115^, including CENP-A ^116^(which appears absent in kinetoplastids) ^117^, modulate levels of centromeric R-loops to limit variation, including genome rearrangement and aneuploidy. Moreover, loss of RAD51 undermines centromere-directed DNA replication initiation in *Leishmania* ^118^. It is also therefore possible that, at least in some circumstances, RAD51 influences R-loop localisation in the absence of an involvement in DNA break repair. Another such circumstance may be seen during transcription initiation. Our mapping data reveal considerable overlap in R-loop localisation in the *rad51*^-/-^ mutants at TSSs relative to H2A.Z localisation in WT cells, suggesting inter- related activities that are normally modulated by RAD51. Such an activity may be independent of RAD51’s role in DNA repair. However, the accumulation of DNA damage marked by yH2A at *T. brucei* TSSs after loss of RNase H2A is intriguing ^38^, as are emerging roles for H2A.Z in DNA repair ^119, 120^ and replication ^121^. BLISS does not predict enrichment of DNA breaks at these sites in *rad51*-/- mutants, though RAD51-dependent R-loops may form due to a lesion, structure or chromatin environment not detected by this approach.

Our data do not reveal a role for RAD51 in TERRA association with telomeres in *T. brucei*. Though this observation differs from mammals and yeast ^71, 72^, it may be because Rad51-directed R-loop formation using TERRA occurs when telomeres are short and use homology-directed repair for maintenance. Previous work has shown that critically short telomeres in *T. brucei* are associated with increased VSG switching ^37, 122, 123^, but R-loop levels and location - or RAD51 involvement - have not been tested. Loss of shelterin components in *T. brucei* has also been shown to activate *VSG* switching ^28, 124, 125^, as has loss of ORC1/CDC6 ^126^, which in mammals associates with TERRA ^127^. It is conceivable that TERRA, acted upon by RAD51, provides an alternative route for *VSG* switching to that we describe here, but where R-loops are also involved.

## Methods

### Trypanosome growth and genetic manipulation

*Trypanosoma brucei* MiTat1.2 (Lister 427) BSF cells and a T7 polymerase/Cas9 expressing derivative line (derived using plasmid pJ1339)^128^ were cultured in HMI-9 medium (Life technologies), supplemented with 10% foetal calf serum at 37 °C with 5% CO2. For epitope tagging and allele knockout, primers were designed with LeishGEdit (http://www.leishgedit.net/) and donor sequences were PCR-amplified from pPOT plasmids (containing mNeonGreen and blasticidin resistance genes) as previously described[64], along with a single guide RNA. Mutant lines were generated by Amaxa transfection with ethanol-purified PCR products and drug-section to generate clonal lines, which were confirmed by PCR.

### Microscopy

For analysis of DNA damage yH2A (1:1,000) antiserum was used, and DNA-RNA hybrids were detected using S9.6 (Kerafast, 1:1,000). To perform immunofluorescence, formaldehyde-fixed parasites were permeabilised with 0.1% IGEPAL and blocked with 1% bovine serum albumin (BSA) for α-yH2A staining. For S9.6 staining, methanol-fixed parasites were instead permeabilised with 0.5% v/v Triton X-100 and blocked with 0.1% w/v BSA plus 0.01% v/v Tween-20 at 37 °C. Alexa Fluor® Plus 488 secondary antibodies were diluted 1:1,000 in the respective blocking solutions before mounting with 5 µL DAPI Fluoromount-G® (Southern Biotech) and imaging. VSG immunofluorescence analysis was performed following the protocol of Glover et al. (2016)^129^ Briefly, formaldehyde fixed parasites were adhered to glass slides and blocked with 50% FCS in PBS for 45 min before staining with primary anti-VSG (1:10,000) and secondary Alexa Fluor (1:1,000) antibodies and mounting with DAPI Fluoromount-G® (Southern Biotech). Imaging was performed with an Axioscope 2 fluorescence microscope (Zeiss) 63x/1.40 oil objective, or Leica DiM8 microscope to acquire Z-stacks of 5-µM thickness in 25 sections. To quantify fluorescence intensity, images were obtained using the same exposure times and were later processed on ImageJ (http://imagej.net/) using the same parameters.

### DRIP-qPCR and DRIP-seq

ChIP-IT® Express Enzymatic (Active Motif, Cat#53035) kit was used for DRIP, as previously described[68]. Briefly, 2 x 10^8^ mid-logarithmic phase cells were fixed with 1% formaldehyde before nuclei were isolated and chromatin digested with an enzymatic cocktail. 1 µg chromatin was used for immunoprecipitation (IP) reactions in 100 µL aliquots with 5 µg S9.6 antibody (Kerafast), as per the kit protocol. On-bead RNase H treatment control was performed for each reaction: washed beads were incubated with 300 µL 1X RNase H Buffer (NEB) plus 30 units of RNase H (NEB) or 15 µL nuclease-free water at 37°C for 3 hr. After chromatin was eluted and reverse crosslinked, DNA was eluted as per the kit protocol. For DRIP-qPCR, eluted DNA and the respective input samples were diluted to 1:10 and 1:100, respectively, before 1 μL was added to a qPCR reaction mix containing 1X Fast SYBR Green Master Mix (Invitrogen) and 400 nM of each forward and reverse qPCR primers. qPCR was performed with a 7500 Real-Time PCR system (Applied Biosystems) as follows: 50 °C for 2 min and 95 °C for 2 min, followed by 40 cycles of 95 °C for 15 sec, 59 °C for 15 sec and 72 °C for 1 min. Fluorescence intensity was measured at the end of the extension step (72 °C for 1 min). Analysis was done by first adjusting the CT values according to the dilution factor, i.e., -log2(dilution 100 factor). Subsequently, the adjusted CT values of the DRIP samples and the inputs were compared. For DRIP-seq, DNA libraries were prepared from the chromatin using a Qiagen FX 96 Library Kit (QIAGEN) for sequencing using the Illumina NextSeq 500 platform, to a depth of 15 million 100-bp paired-end reads per sample.

### Long-read de novo genome assembly

For long-read assembly, DNA of *Trypanosoma brucei* MiTat1.2 (Lister 427) BSF cells was extracted using MagAttract HMW DNA kit (Qiagen) and sequenced using an Oxford Nanopore Technologies (ONT) MinION device with R9.4.1 chemistry. Basecalling was done using guppy (Linux CPU version 3.3.3). A total of 319075 reads (N50 28.46 kb) were generated and subsequently used for genome assembly using Canu (default settings, predicted genome size = 35 Mb). The resulting assembly was polished with four iterations of Pilon using paired-end (2x75bp) Illumina data from the same strain of parasites.

### DRIP-seq data analysis

Analysis was performed as previously described ^66^. Briefly, TrimGalore was used to trim reads (Phred Score <20) before mapping to either the *T. brucei* Lister 427 v10 ^69^ or Oxford Nanopore assembly genome (this work) using Bowtie2 in “very-sensitive” mode. Mapped reads with MapQ value < 1 were filtered using SAMtools and the fold-change between the read depths of DRIP IP DNA relative to input control was calculated in 50-bp bins using DeepTools bamCompare tool, where background was separated from DRIP signals using signal extraction scaling. deepTools computeMatrix, plotProfile, and plotHeatmap functions were used to generate plots.

### Breaks labelling in situ and sequencing (BLISS)

BLISS was performed by adapting the the protocol developed by Bouwman et al^73^, summarised here. Cells were fixed in 2% methanol-free formaldehyde for 10 min at room temperature, before quenching with 125 μM glycine. 1 x 10^8^ cross-linked cells were then lysed using 200 µL lysis buffer 1 (10 mM Tris-HCl, 10 mM NaCl, 1 mM EDTA, 0.2% Triton X-100, pH 8 at 4 °C) and resuspended in 200 µL of pre-warmed lysis buffer 2 (10 mM Tris-HCl, 150 mM NaCl, 1 mM EDTA, 0.3% SDS, pH 8 at RT). DSBs were blunted in situ using the Quick Blunting Kit (NEB) and BLISS linkers were ligated overnight at 16 °C using T4 DNA ligase (ThermoFisher). The nuclei were washed with 200 µL 1X CutSmart Buffer (NEB) supplemented with 0.1% Triton X-100 to remove un-ligated linkers before being resuspended in TAIL buffer (10 mM Tris, 100 mM NaCl, 50 mM EDTA, 1% SDS, pH 7.5 at RT), with added Proteinase K (NEB). Samples were incubated, shaking overnight at 55 °C before DNA was extracted using a phenol/chloroform/isoamyl alcohol (PCI) mixture and ethanol precipitated into 100 µL TE buffer. Sonication with a BioRuptor (30 cycles of 30 s on, 60 s off at high intensity) sheared the DNA which was then concentrated using AMPure XP beads and quantified using Qubit dsDNA BR assay (ThermoFisher Scientific). To prepare libraries, in vitro transcription (IVT) of 100-200 ng of BLISS template gDNA was performed in DNA LoBind tubes using MEGAscript T7 (ThermoFisher) at 37 °C for 14-15 h (lid temperature set at 70 °C). The DNA template was subsequently degraded using 2U DNase I (ThermoFisher) and RNA was purified using pre-warmed RNAClean XP beads (Beckman Coulter) at room temperature. Small RNA Seq 3’ adapters (Illumina) were ligated using T4 RNA ligase (NEB) at 25 °C for 2 h, and reverse transcription was performed with SuperScript IV reverse transcriptase (ThermoFisher) at 50 °C for 50 mins and stopped at 80 °C for 10 mins. The resultant library was amplified using NEBNext Ultra II Q5 Master Mix by performing 8 cycles initially, then bead-purifying the mixture using 0.8 AMPure XP DNA beads, before further amplification via 4 additional cycles of PCR. Finally, DNA libraries were purified using two-sided AMPure purification and sequenced on an Illumina NextSeq 500 platform to a depth of 60 million single-end reads per sample. Sequencing was done at Glasgow Polyomics.

BLISS reads were trimmed to remove library barcodes and UMIs using cutadapt, allowing up to 1 mismatch, and the library barcode and UMI were added to the read name using a custom python script, cutadaptInfoToFastq.py. Reads were then mapped to the *Trypanosoma brucei* Lister 427 v10 genome using bowtie2 under default settings. The aligned reads were exported in BAM format before sorting with SAMtools and deduplicating using the UMI with umi_tools dedup. This ensured that reads mapping to the same locations with different UMIs come from distinct DSBs rather than being PCR duplicates. Finally, the filtered reads were normalized with bamCoverage and then exported as a bigwig file. The bigwig files were either visualized as tracks using Integrated Genome Viewer or plotted as signal profiles or heatmaps as described for DRIP-seq.

Read depth peaks were then detected as a proxy for regions of DNA breaks using the macs2 peak caller version 2.2.7.1^74^. We first filtered the alignment bam files to retain only reads that could be confidently assigned to a single genomic region by requiring reads to have mapping quality of at least 10 and excluded alignments marked as “supplementary” by the aligner (bwa mem). For peak calling we used “macs2 callpeak” with these settings: effective genome size (-g) 35000000, keep all duplicates (--keep-dup all), do not build model (--nomodel), extend reads by 80 bp (--extsize 80). Other parameters were set to default to select only peaks with FDR < 0.05, and without an input control. The analysis pipeline of the BLISS data is available at https://github.com/glaParaBio/bliss_dna_break_analysis as Snakemake pipeline.

## Data availability

BLISS and DRIP-seq sequences are available at the NCBI Sequence Read Archive under project number PRJEB61713. Nanopore and Illumina reads used for genome assembly have been deposited to the NCBI Sequence Read Archive (SRA) under project number PRJNA962304. DRIP-MS data and availability is described in Girasol et al, BIORXIV/2023/540366..

## Acknowledgements

This work was funded by the Philippine Council for Health Research and Development (DOST-PCHRD) for M.G., Wellcome Trust (Investigator Award (224501/Z/21/Z) to R.M., Henry Wellcome fellowship (218648/Z/19/Z) to E.M.B., and an Institutional Strategic Support Fund (ISSF3) award held at the University of Glasgow (204820/Z/16/Z) to E.M.B. and R.M.), and by the BBSRC (BB/N016165/1 and BB/R017166/1 to R.M, and to BB/W001101/1 to R.M. and C.A.M). The Wellcome Centre for Integrative Parasitology is funded by a Wellcome Trust Strategic Award (104111/Z/14/Z/A). We thank all staff at Glasgow Polyomics for DNA sequencing advice and generation.

## Supplementary figure legends

**Figure S1.**
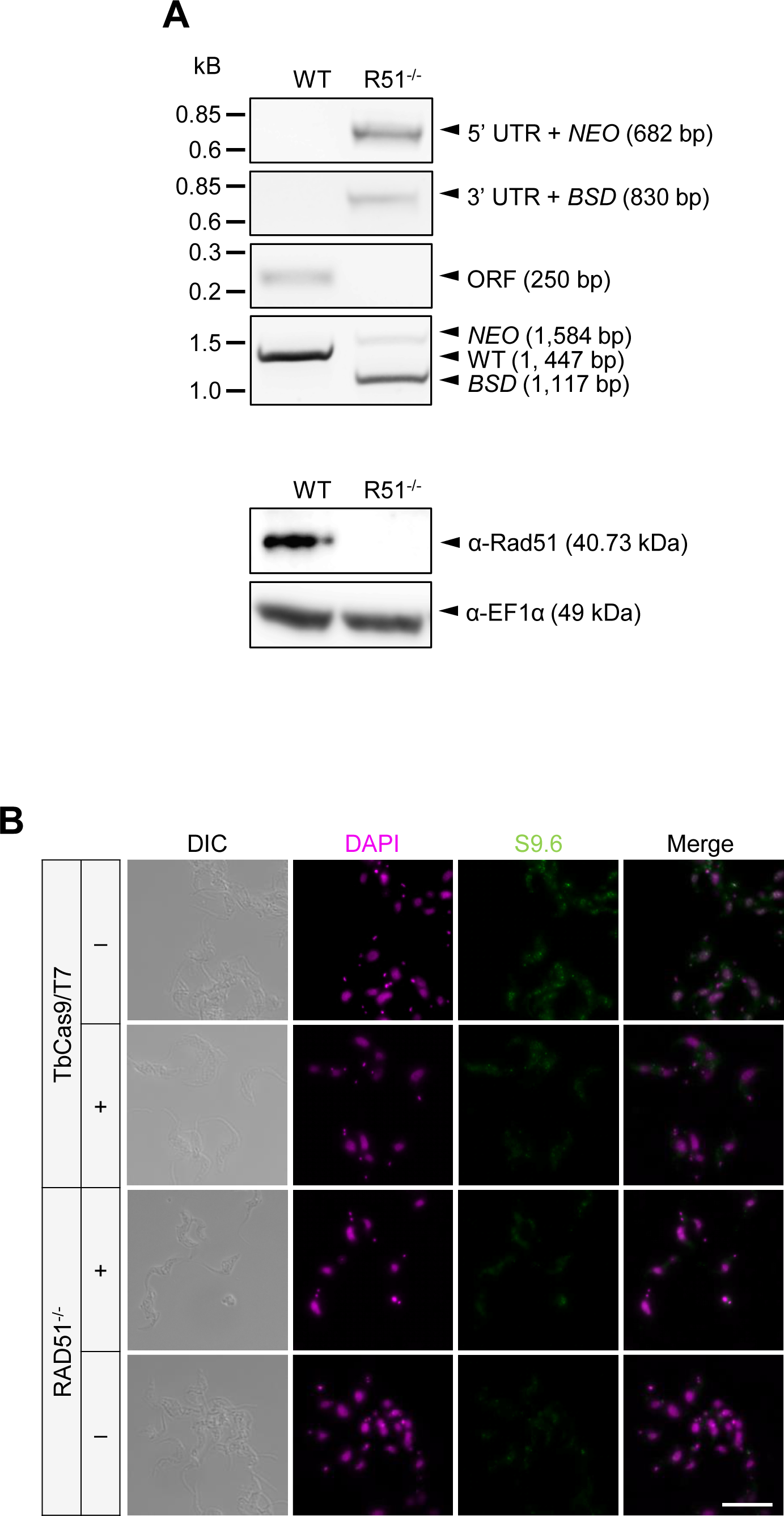
CRISPR-Cas9 generation and analysis of *T. brucei RAD51* null mutants. (A) Upper agarose gels show PCR of a *rad51*-/- transformant, detailing the expected integration of NEO and BSD resistance cassettes, replacing the *RAD51* ORF; P, parental TbCas9/T7 (‘wild type’) cell line. Lower gel shows a western blot of RAD51 protein expression in the same mutant using an anti-RAD51 antibody; EF1α was used as a loading control. (B) Sample images of S9.6 immunofluorescence, comparing parental TbCas9/T7 cells and *rad51*-/- mutants, with (+) or without (-) *E. coli* RNase HI treatment; scale bar = 10 µm.

**Figure S2.**
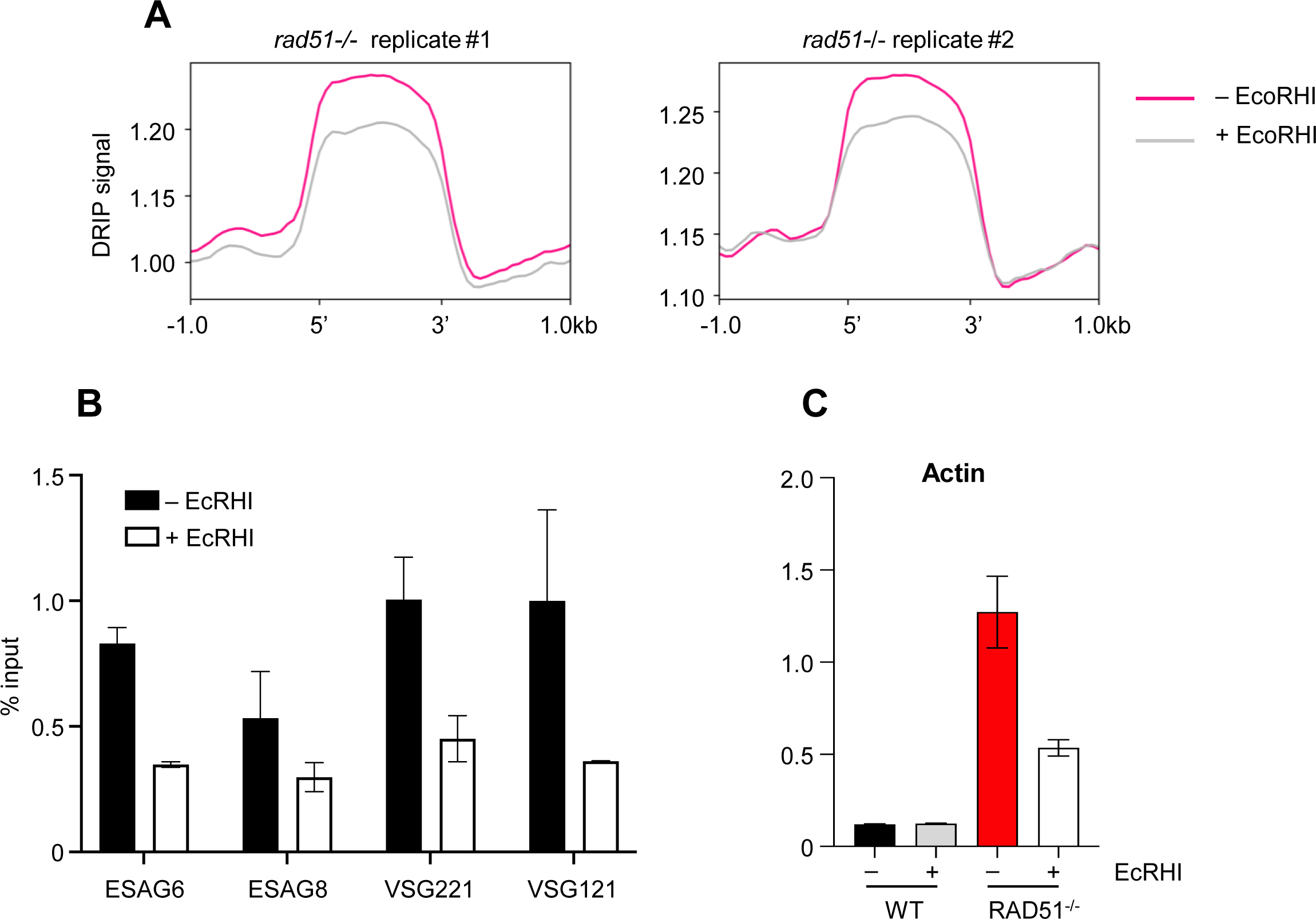
Testing DRIP efficacy. (A). Analysis of DRIP-seq data from two independent *rad51*-/- mutants in the presence (grey) or absence (magenta) of *E. coli* RNase HI; graphs depict metaplots of reads mapped to all RNA Pol II–transcribed CDSs (scaled to size), including +/- 1 kb. (B) DRIP-qPCR of *rad51*-/- cells, quantifying recovery of R-loops from BES1 (*ESAG6*, *ESAG8*, *VSG221*) and BES3 (*VSG121*). (C) DRIP-qPCR comparing wild type and *rad51-*/- R-loop enrichment within the actin CDS. In all DRIP-qPCR analyses, abundance is expressed as a percentage in IP relative to input sample, with or without treatment with *E. coli* RNase HI; error bars represent SEM of two independent replicates.

**Figure S3.**
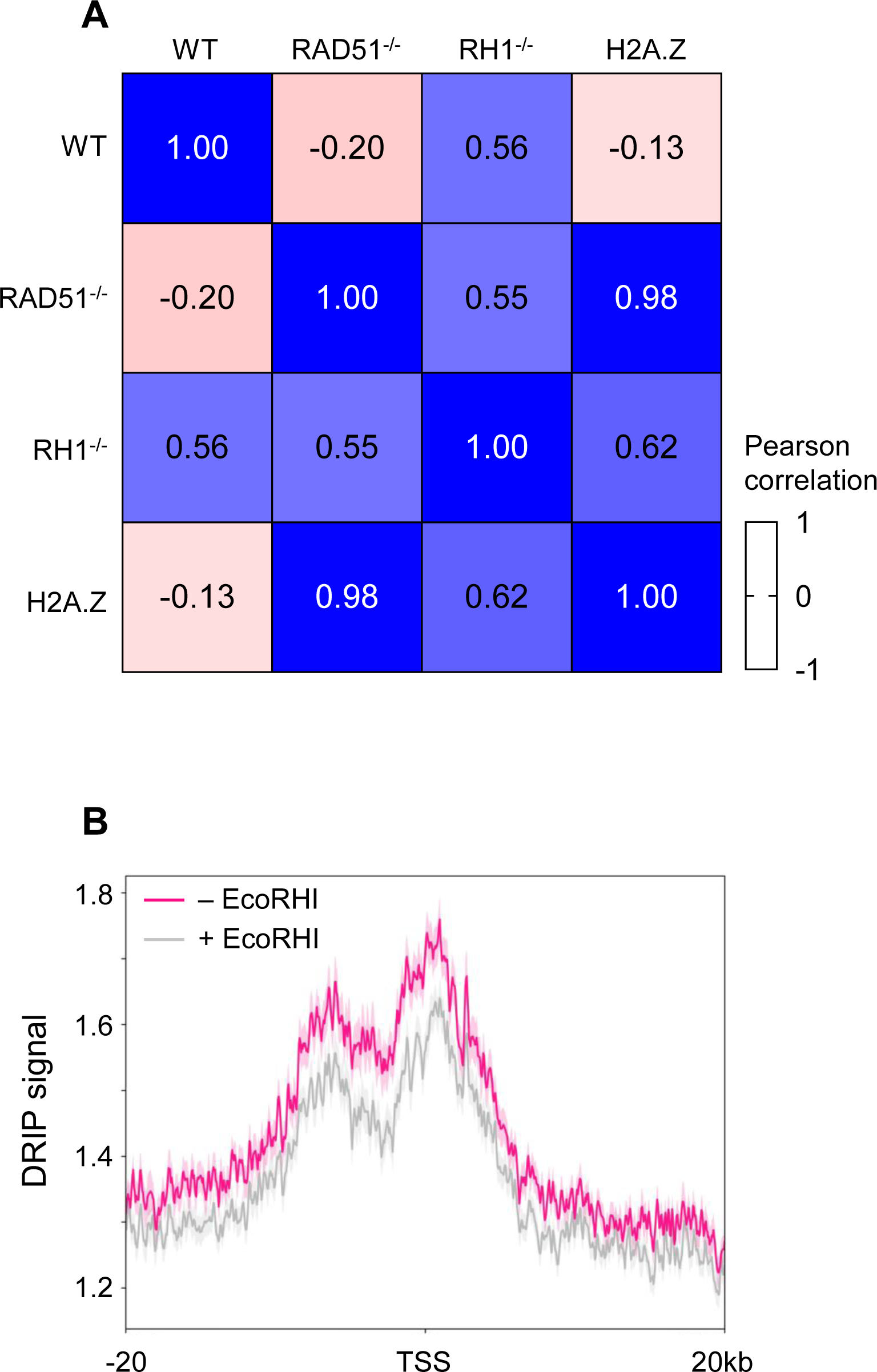
DRIP-seq signal correlates with H2A.Z enrichment in transcription start sites in *rad51-/-* mutants. (A) Pearson correlation coefficients of DRIP-seq signal in wild-type (WT), *rad51-*/- and *RNase H1*-/- (*RH1*-/-) cells relative to H2A.Z ChIP-seq signal in WT cells across a 40-Kb genomic window surrounding all TSSs. (B) Metaplots of *rad51-/-* DRIP-seq reads along a 40-Kb window surrounding the H2A.Z ChIP-seq peaks at TSSs, showing the difference between samples treated and not treated with *E. coli* RNase HI.

**Figure S4.**
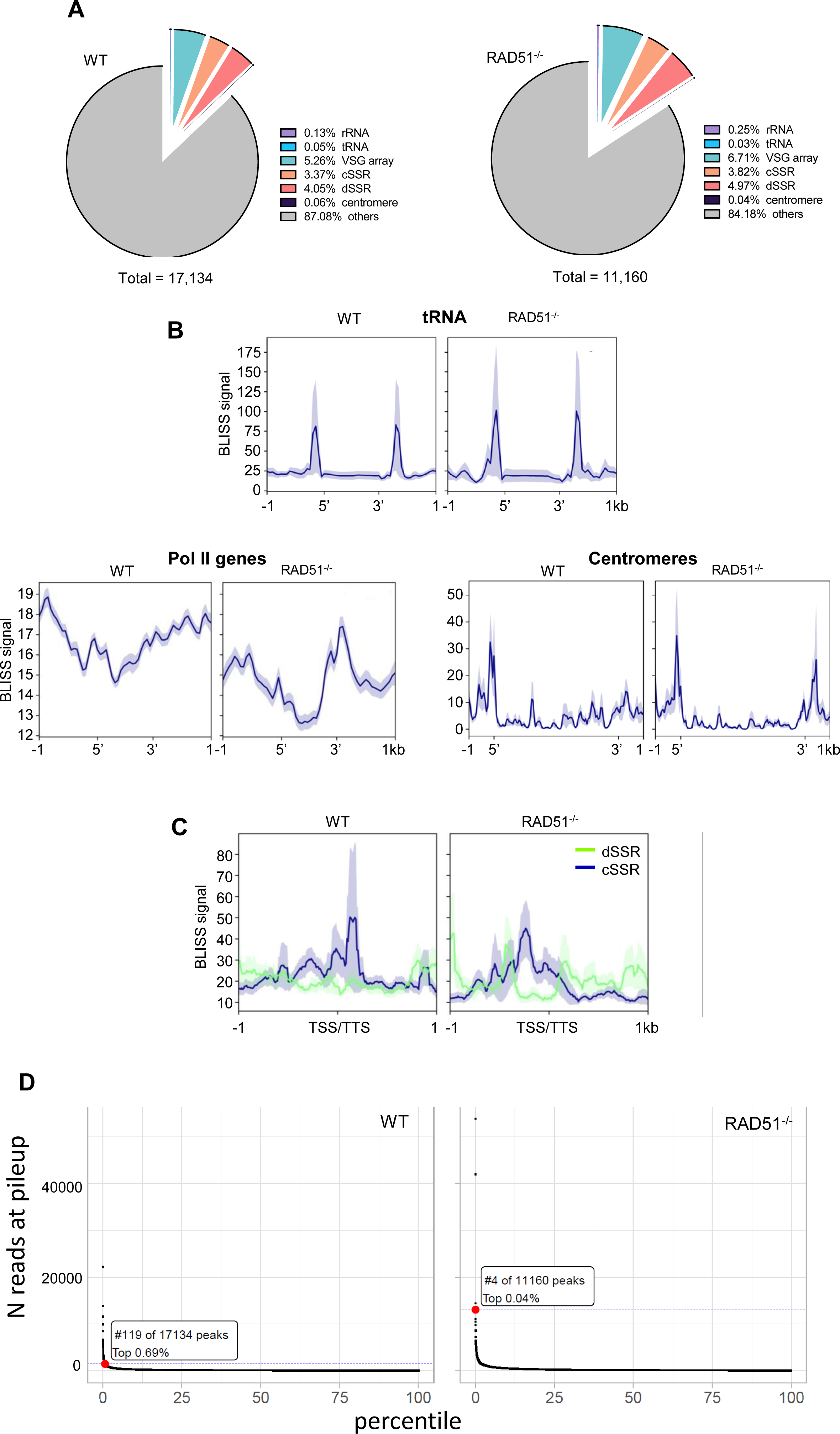
Genome-wide analysis of DNA breaks in *T. brucei* BSF cells by BLISS. (A) Distribution of MACS2- predicted peaks of BLISS signal in wild type (WT) and *rad51*-/- cells; proportions of total BLISS peaks residing in different genomic locations are shown; ‘others’ represents RNA Pol II PTUs. (B) Ranking of BLISS peak signal in WT and *rad51-/-* cells; N reads at pileup represents BLISS reads/million reads at the summit of a BLISS peak; BLISS reads mapping to active BES1 are highlighted as a red dot. (C) Mapping of BLISS signal (read count per million mapped reads) across selected genome features (data are shown as average normalised signal across the region of interest); in each case, metaplots depict the mean signal plus SEM (shaded) across all CDSs or centromeres (scaled to the same size), including 1 Kb of upstream and downstream sequence. (D) Metaplots comparing BLISS signal around all predicted TSSs or TSSs (identified as divergent (d) or convergent (c) strand switch regions, SSRs), in wildtype (WT) and *rad51*-/- cells.

**Figure S5.**
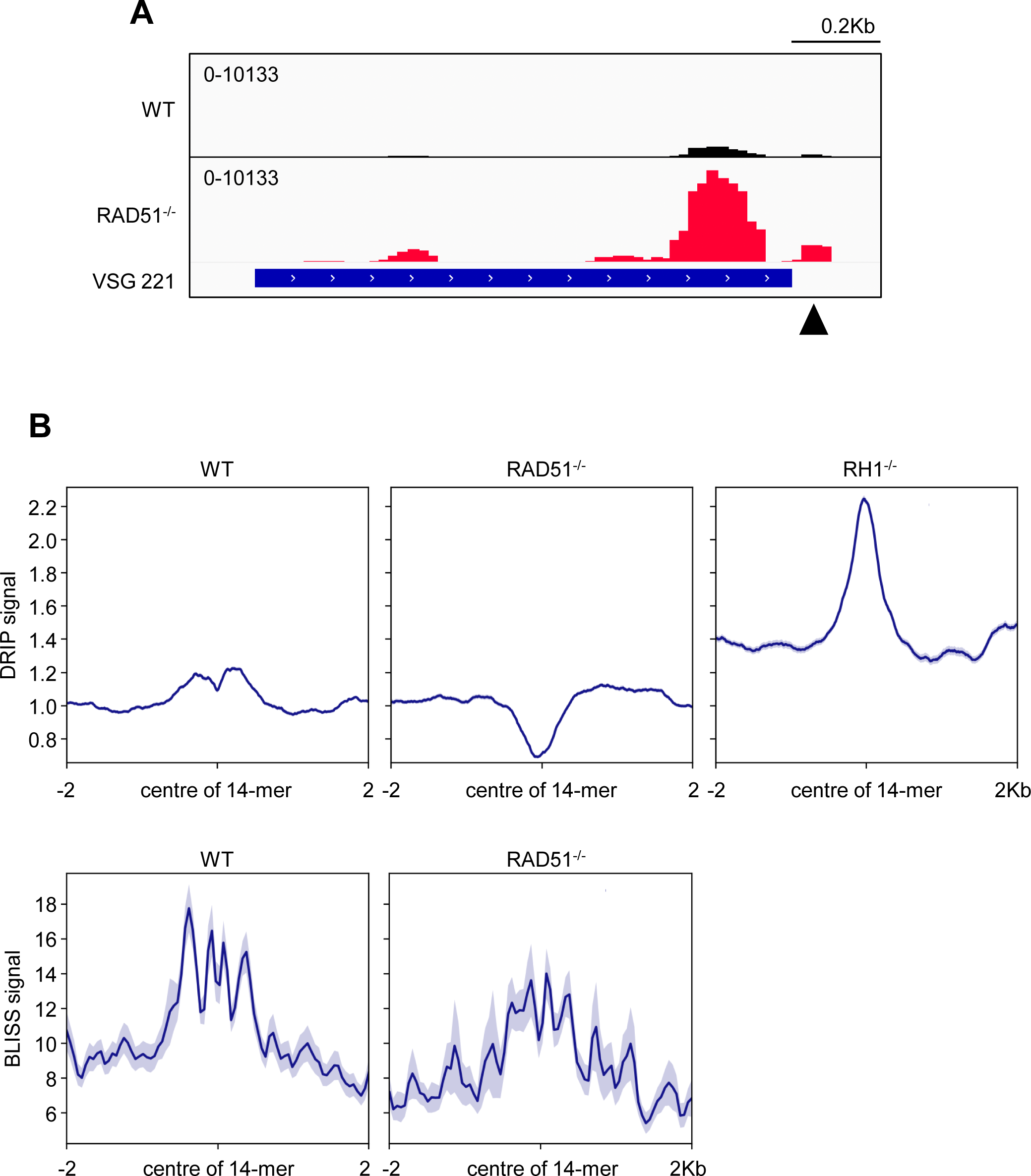
DSB and R-loop distribution around the conserved *VSG* 14-mer motif. (A) IGV track showing BLISS signal (read counts per million mapped reads) around the actively expressed *VSG* gene, *VSG221*; arrow denotes the location of the conserved 14-mer sequence. (B) Metaplot showing distribution of DRIP-seq signal (DRIP/input) in sequences centred on the 14-mer at the 3’ end of all *VSG* genes, comparing wild type, *rad51*-/- and *RNase H1*-/- cells. (C) Metaplot showing BLISS signal (mean reads per million mapped reads, and SEM shaded) centred on the 14-mer of all *VSG*s, comparing wild type and *rad51*-/- cells.

## Notes

### Competing Interest Statement

The authors have declared no competing interest.

